# A sulfoglycolytic Entner-Doudoroff pathway in *Rhizobium leguminosarum* bv. *trifolii* SRDI565

**DOI:** 10.1101/2019.12.17.868638

**Authors:** Jinling Li, Ruwan Epa, Nichollas E. Scott, Dominic Skoneczny, Mahima Sharma, Alexander J.D. Snow, James P. Lingford, Ethan D. Goddard-Borger, Gideon J. Davies, Malcolm J. McConville, Spencer J. Williams

## Abstract

Rhizobia are nitrogen fixing bacteria that engage in symbiotic relationships with plant hosts but can also persist as free-living bacteria with the soil and rhizosphere. Here we show that free living *Rhizobium leguminosarum* SRDI565 can grow on the sulfosugar sulfoquinovose (SQ) using a sulfoglycolytic Entner-Doudoroff (sulfo-ED) pathway resulting in production of sulfolactate (SL) as the major metabolic end-product. Comparative proteomics supports the involvement of a sulfo-ED operon encoding an ABC transporter cassette, sulfo-ED enzymes and an SL exporter. Consistent with an oligotrophic lifestyle, proteomics data revealed little change in expression of the sulfo-ED proteins during growth on SQ versus mannitol, a result confirmed through biochemical assay of sulfoquinovosidase activity in cell lysates (data are available via ProteomeXchange with identifier PXD015822). Metabolomics analysis showed that growth on SQ involves gluconeogenesis to satisfy metabolic requirements for glucose-6-phosphate and fructose-6-phosphate. Metabolomics analysis also revealed the unexpected production of small amounts of sulfofructose and 2,3-dihydroxypropanesulfonate, which are proposed to arise from promiscuous activities of the glycolytic enzyme phosphoglucose isomerase and a non-specific aldehyde reductase, respectively. This work shows that rhizobial metabolism of the abundant sulfosugar SQ may contribute to persistence of the bacteria in the soil and to mobilization of sulfur in the pedosphere.

## Introduction

Sulfur is essential for plant growth and is the fourth most important macronutrient after nitrogen, phosphorus, and potassium. Up to 10 kg/ha/y of sulfur is deposited in rain, especially near industrialized areas.^1^ However, sulfur dioxide emissions from industrial sources have decreased in recent decades as a result of pollution mitigation and the move to low sulfur fuels and renewable energy sources, and quantities received from atmospheric sources is now at levels below that required by most crops.^2^ Sulfur deficiency in soils is primarily combated by application of sulfur-containing fertilizers such as superphosphate, ammonium sulfate and gypsum,^3^ which are applied across all major cropping and pasture areas worldwide.^4^ Soils contain significant amount of sulfur, yet plants can only use sulfur in the form of sulfate and it has been shown that 95-98% of sulfur in soils is in the form of unavailable biological sulfur.^4^ Thus, effective microbial cycling of sulfur from biological to inorganic forms within the soil is important^5^ and has the potential to enhance crop yields and reduce reliance on fertilizers.

It is estimated that around 10 billion tonnes per annum of the sulfosugar sulfoquinovose (SQ) is produced annually by photosynthetic organisms, including plants, cyanobacteria and algae.^6^ SQ is primarily found as the glycerolipid sulfoquinovosyl diacylglycerol (SQDG), and land plants can contain as much as 10% SQDG in their thylakoid membrane glycerolipids.^7^ Very little is known about how SQ is metabolized within soils, although it has been shown to undergo very rapid mineralization to inorganic sulfate.^8^ based on X-ray absorption near-edge spectroscopy measurements, it is estimated that 40% of sulfur within various sediments and humic substances exist as sulfonate.^9^

Bacteria are likely to be primarily responsible for the biomineralization of SQ, possibly by using SQ as a carbon source and catabolizing it via a modified version of glycolysis, termed sulfoglycolysis.^10^ Two sulfoglycolytic processes have been described: the sulfoglycolytic Embden-Meyerhof-Parnas (sulfo-EMP) pathway,^11^ and the sulfoglycolytic Entner-Doudoroff (sulfo-ED) pathway (Fig. 1).^12^ The sulfo-ED pathway was first reported in *Pseudomonas putida* strain SQ1, a bacterium isolated from freshwater sediment, catabolised of SQ with excretion of equimolar amounts of sulfolactate (SL).^12^ The sulfo-ED operon of *P. putida* SQ1 contains 10 genes including a transcriptional regulator, an SQ importer and SL exporter, a sulfoquinovosidase, SQ mutarotase, SQ dehydrogenase, SL lactonase, SG dehydratase, KDSG aldolase and SLA dehydrogenase enzymes. Based on genome-wide annotation studies, the sulfo-ED pathway is predicted to occur in a range of alpha-, beta- and gamma-proteobacteria.^12^ However, no direct evidence for this pathway has been reported for any organism other than *P. putida* SQ1. Other members of the microbial community can catabolize SL and 2,3-dihydroxypropanesulfonate (DHPS; the product of the sulfo-EMP pathway) to inorganic sulfur,^13^ completing the biomineralization of SQ.

**Figure 1:**
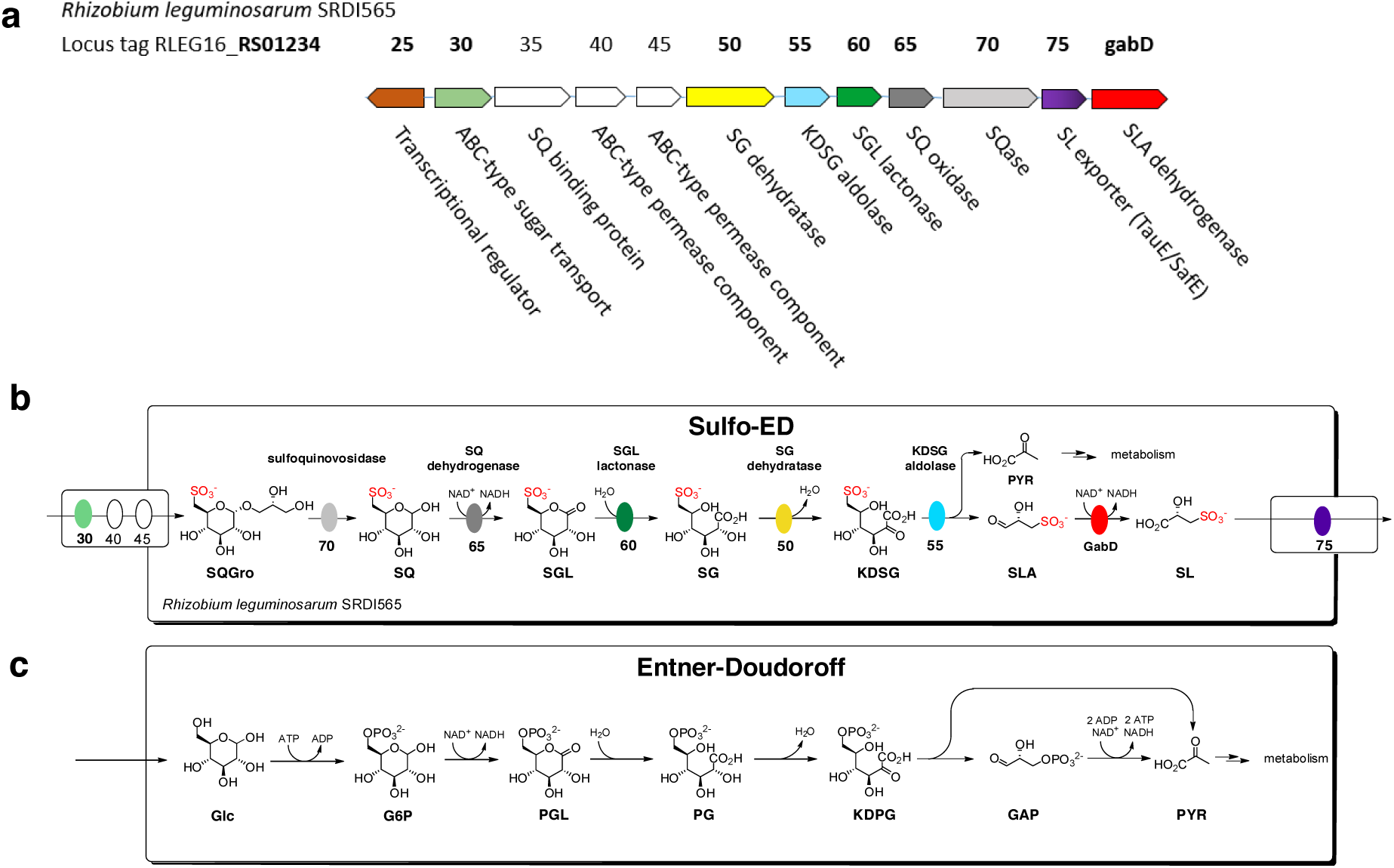
Proposed sulfoglycolytic Entner-Doudoroff (sulfo-ED) pathway in *Rhizobium leguminosarum* bv. t*rifolii* SRDI565. (a) Operon encoding sulfo-ED pathway. (b) Proposed sulfo-ED pathway. (c) Comparison with the Entner-Doudoroff pathway.

*Rhizobium leguminosarum* bv. *trifolii* SRDI565 (syn. N8-J), hereafter *Rl*-SRDI565, was isolated from a soil sample collected in western New South Wales but has the capacity to colonize *Trifolium subterraneum* subsp. *subterraneum* (subterranean clover) and other *Trifolium* spp.^14^ Colonization of trifolium species with *Rl*-SRDI565 results in suboptimal nodulation and nitrogen fixation in some species and ineffective nitrogen fixation in others, leading to reduced shoot nitrogen content relative to other commercial strains.^15^ Interestingly, the genome of *Rl*-SRDI565 encodes all the genes needed for a functional sulfo-ED pathway,^16^ although there is no evidence to show that this is operative and/or that *Rl*-SRDI565 can use SQ as a major carbon source.

Rhizobia participate in sophisticated symbiotic relationships with leguminous host plants that allow them to fix atmospheric dinitrogen to provide a growth advantage to the host.^17^ Symbiosis is triggered by molecular communication between the bacterium and the host resulting in nodule formation on the root and colonization by the bacterium. Within nodule bacteroids the energy intensive fixation of nitrogen is supported by C_4_-dicarboxylates (primarily malate, fumarate, and succinate) sourced from glycolysis of sucrose photosynthate within the plant host.^17^ Owing to the importance of biological nitrogen fixation for input of nitrogen into the biosphere, the symbiosis of rhizobia and leguminous hosts has been well studied. However, rhizobia can also exist as free-living bacteria within the soil and rhizosphere.^18^ Here, like other soil bacteria, they adopt a saprophytic and oligotrophic lifestyle where they utilize a variety of alternative carbon sources, including a wide range of carbohydrates.^19^ Most likely, the ability of various rhizobia to persist in the pedosphere depends upon their ability to utilize diverse carbohydrate and non-carbohydrate substrates and establish an appropriate niche. SQ or its glycosides are likely to be a common soil constituent and nutrient given its ubiquitous production by plants. Possibly, the sulfo-ED pathway in *Rl*-SRDI565 might provide it with the capacity to survive on plant derived SQ or SQDG in the rhizosphere and in the soil.

Here we investigated whether the sulfo-ED pathway is active in *Rl*-SRDI565 and its potential role in utilizing plant-derived SQ or SQDG in the rhizosphere and in the soil. We show that *Rl*-SRDI565 can grow on SQ and sulfoquinovosyl glycerol (SQGro) as sole carbon source. Growth on SQ leads to excretion of SL into the growth media indicating active sulfoglycolysis. This was supported by proteomic analyses, which showed that several genes in the sulfo-ED operon are upregulated when bacteria are grown on SQ, while metabolomic analyses confirm the presence of characteristic intermediates of the sulfo-ED pathway, as well as the unexpected production of intracellular DHPS. Overall, we show that *Rl*-SRDI565 has an active pathway for SQ utilization which may support growth of this bacterium in the environment, and in turn provides a new model organism for the study of the sulfo-ED pathway.

## Results

Analysis of the genome of *Rl*-SRDI565 revealed a sulfo-ED operon that had the same genes, but no synteny with the *P. putida* SQ1 operon (Figure 1). Genes with high sequence identity to the *P. putida* proteins included a putative SQase, SQ dehydrogenase, SL lactonase, SG dehydratase, KDSG aldolase and SLA dehydrogenase, and an SL exporter (see Figures S1-S6). The *Rl*-SRDI565 operon contains some important differences compared to that of *P. putida* SQ1. In particular, it lacks a putative SQ mutarotase,^20^ and appears to use an ABC transporter cassette to import SQ/SQGro in place of an SQ/SQGro importer/permease. The putative sulfo-ED pathway in *Rl*-SRDI565 is consistent with the proposed protein functions outlined in Figure 1b, with a comparison to the classical ED pathway in Figure 1c.

Initial attempts were made to grow *Rl*-SRDI565 in completely defined medium, such as M9 minimal media containing 125 μg mL^−1^ biotin ^21^, to allow assessment of different carbon sources on bacterial growth. However, optimal growth could only be achieved using a yeast extract-based medium ^15^. In particular robust growth was achieved using a 5% dilution of 1 g L^−1^ yeast extract (Y_5%_ media) containing 5 mmol mannitol (Y_5%_M), while no detectable bacterial growth was observed on Y_5%_ media alone. Significantly, *Rl*-SRDI565 also grew robustly on Y_5%_ media containing 5 mM SQ (Y_5%_SQ) and reached the same final OD_600_ value as in Y_5%_M (Figure 2a). *Rl*-SRDI565 also grew on Y_5%_ media containing glucose, although to a lower final OD_600_ than in Y_5%_M or Y_5%_SQ. ^13^C NMR spectroscopic analysis of the culture media of stationary phase *Rl*-SRDI565 grown in Y_5%_SQ revealed the presence of three major signals corresponding to SL (Figure 2b). A fourth signal was also observed but not assigned and was also present in stationary phase media of cells grown on Y_5%_M, suggesting it is derived from other carbon sources in the yeast extract. *Rl*-SRDI565 also grew on Y_5%_ containing SQGro, but less robustly than on SQ.

**Figure 2:**
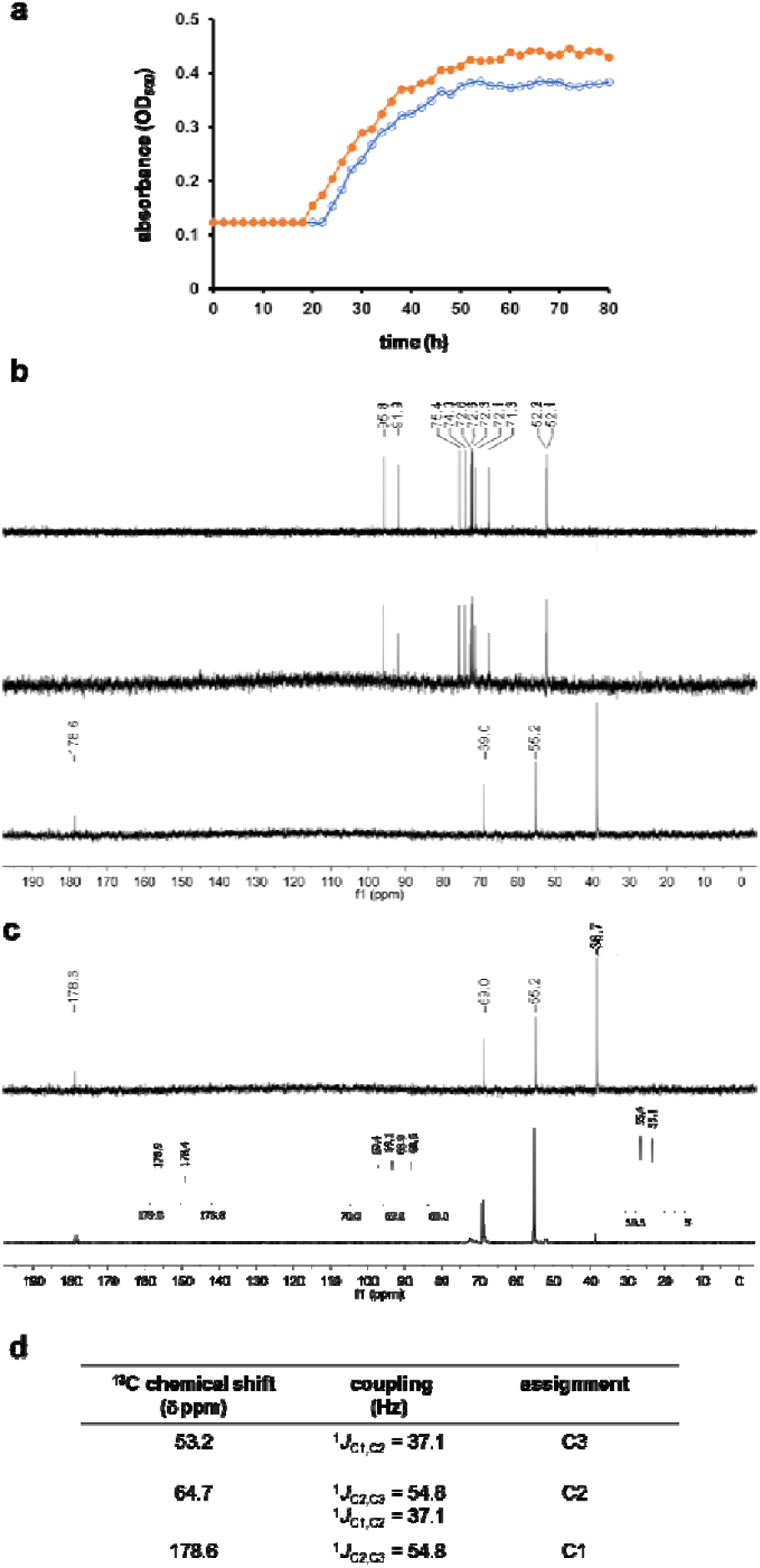
Growth of *Rhizobium leguminosarum* bv. t*rifolii* SRDI565 on SQ produces SL as the major terminal metabolite. a) Growth of *Rl*-SRDI565 on 5% yeast extract media containing 5 mM SQ (●) or 5 mM mannitol (○). This data is representative of 2 independent experiments. b) ^13^C NMR (126 MHz) spectra of (top) SQ, (middle) 5 mM SQ in 5% yeast extract media and (bottom) spent culture media from growth of *Rl*-SRDI565 on 5 mM SQ. c)^13^C NMR (126 MHz) spectrum of spent culture media from growth of *Rl*-SRDI565 on 5 mM (^13^C_6_)-SQ. The signal at δ 38.7 ppm is present in control experiments of *Rl*-SRDI565 grown on mannitol and is believed to derive from yeast extract. d) Tabulated ^13^C NMR (126 MHz) data for ^13^C_3_-SL from (c). All samples contain 10% D_2_O, added to allow frequency lock.

We next examined changes in the proteome of *Rl*-SRDI565 cultivated on mannitol versus SQ. Label-free based quantitative proteomic analysis of five experimental replicates of *Rl*-SRDI565 cultivated on each carbon source, identified 2954 proteins, with 1943 proteins quantified in at least 3 experimental replicates under each growth condition (Supplementary Table 1). Expression levels of 30 proteins potentially associated with SQ metabolism were significantly elevated (-log_10_(p)>2 and a fold change greater than 2 log_2_ units) in bacteria cultivated in Y_5%_SQ (Figures 3a and b). In particular, a suspected KDSG aldolase (annotated as alpha-dehydro-beta-deoxy-D-glucarate aldolase, WP_017967308.1), a member of the proposed sulfo-ED pathway, was significantly increased (-log_10_(p)= 4.74429 and a fold change of 2.38 log_2_). Consistent with the involvement of this pathway we also observed a significant yet less dramatic increase in the proposed SQase (annotated alpha-glucosidase, WP_017967311.1) (-log_10_(p)= 1.43643 and a fold change of 1.02 log_2_). Additional members of the predicted pathway expressed at higher levels in SQ-fed bacteria included the suspected SQ dehydrogenase (annotated as SDR family oxidoreductase, WP_017967310.1) identified by MS/MS events in 4 out of 6 SQ experiments compared to 1 mannitol experiment and the suspected SG dehydratase (annotated as dihydroxy-acid dehydratase, WP_017967307.1) identified by MS/MS events in 3 out of 6 SQ experiments compared to 0 mannitol experiments (Figure S7). However, owing to their low abundance they could not be accurately quantified (Figure S8).

**Figure 3:**
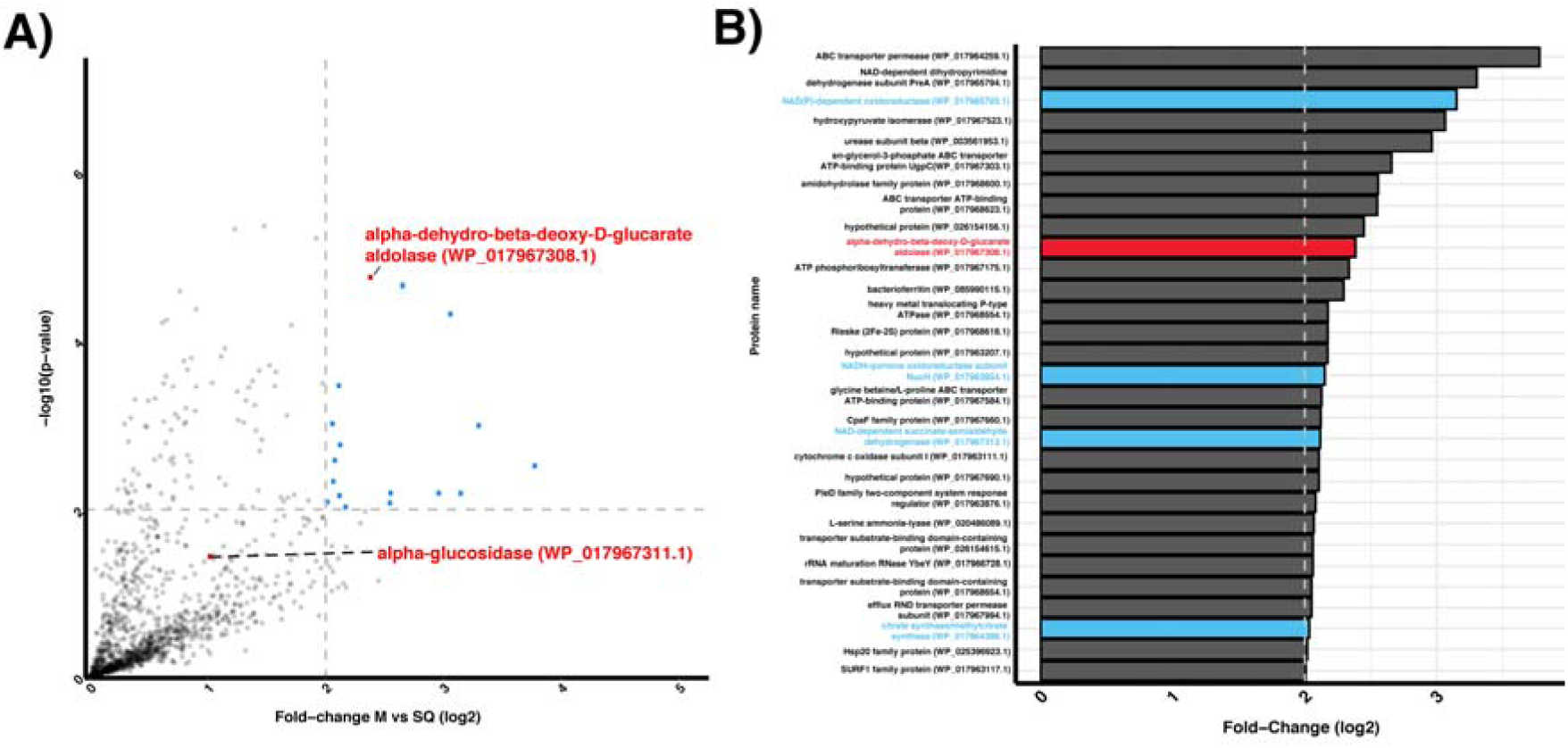
Proteomic analysis of *Rhizobium leguminosarum* SRDI565 growth in sulfoquinovose. Quantitative proteomics was undertaken to identify proteins associated with sulfoquinovose catabolism versus mannitol. A) Examination of proteins observed to increase in abundance greater than four-fold revealed 30 proteins including alpha-dehydro-beta-deoxy-D-glucarate aldolase (WP_017967308.1). B) Growth in sulfoquinovose leads to the increase of multiple proteins associated with the TCA cycle including NAD(P)-dependent oxidoreductase (WP_017965793.1), NADH-quinone oxidoreductase subunit NuoH (WP_017963854.1), NAD-dependent succinate-semialdehyde dehydrogenase (WP_017967313.1) and citrate synthase/methylcitrate synthase (WP_017964386.1) highlighted in blue.

Other proteins that were significantly increased in SQ-fed bacteria included a NAD(P)-dependent oxidoreductase (WP_017965793.1), NADH-quinone oxidoreductase subunit NuoH (WP_017963854.1), a NAD-dependent succinate-semialdehyde dehydrogenase (WP_017967313.1) and a citrate synthase/methylcitrate synthase (WP_017964386.1) supporting an alteration with the TCA cycle and oxidative phosphorylation under conditions of growth on SQ (Figure 3b).

To demonstrate activity for a representative sulfo-ED enzyme from *Rl*-SRDI565, we cloned and expressed the gene encoding the putative SQase. To support future structural studies, we expressed the N-terminal hexahistidine tagged K375A/K376A variant, termed *Rl*SQase*, a mutant enzyme whose design was guided by the Surface Entropy Reduction prediction (SERp) server (Figure S9).^22^ Size exclusion chromatography-multiple angle light scattering (SEC-MALS) analysis of *Rl*SQase* revealed that the protein exists as a dimer in solution (Figure S10). Enzyme kinetics were performed using the chromogenic SQase substrate 4-nitrophenyl α-sulfoquinovoside (PNPSQ). *Rl*SQase* exhibited a bell-shaped pH profile with optimum at pH 7-8 and consistent with titration of catalytically important residues of pKa1 = 6.5 ± 0.4 and pKa2 = 8.6 ± 0.3. The enzyme displayed saturation kinetics with Michaelis-Menten parameters *k*_cat_ = 1.08 ± 0.17 s^−1^, *K*_M_ = 0.68 ± 0.25 mM, and *k*_cat_/*K*_M_ = 1.59 ± 0.83 s^−1^ mM^−1^ (Figure 4a and 4b).

**Figure 4:**
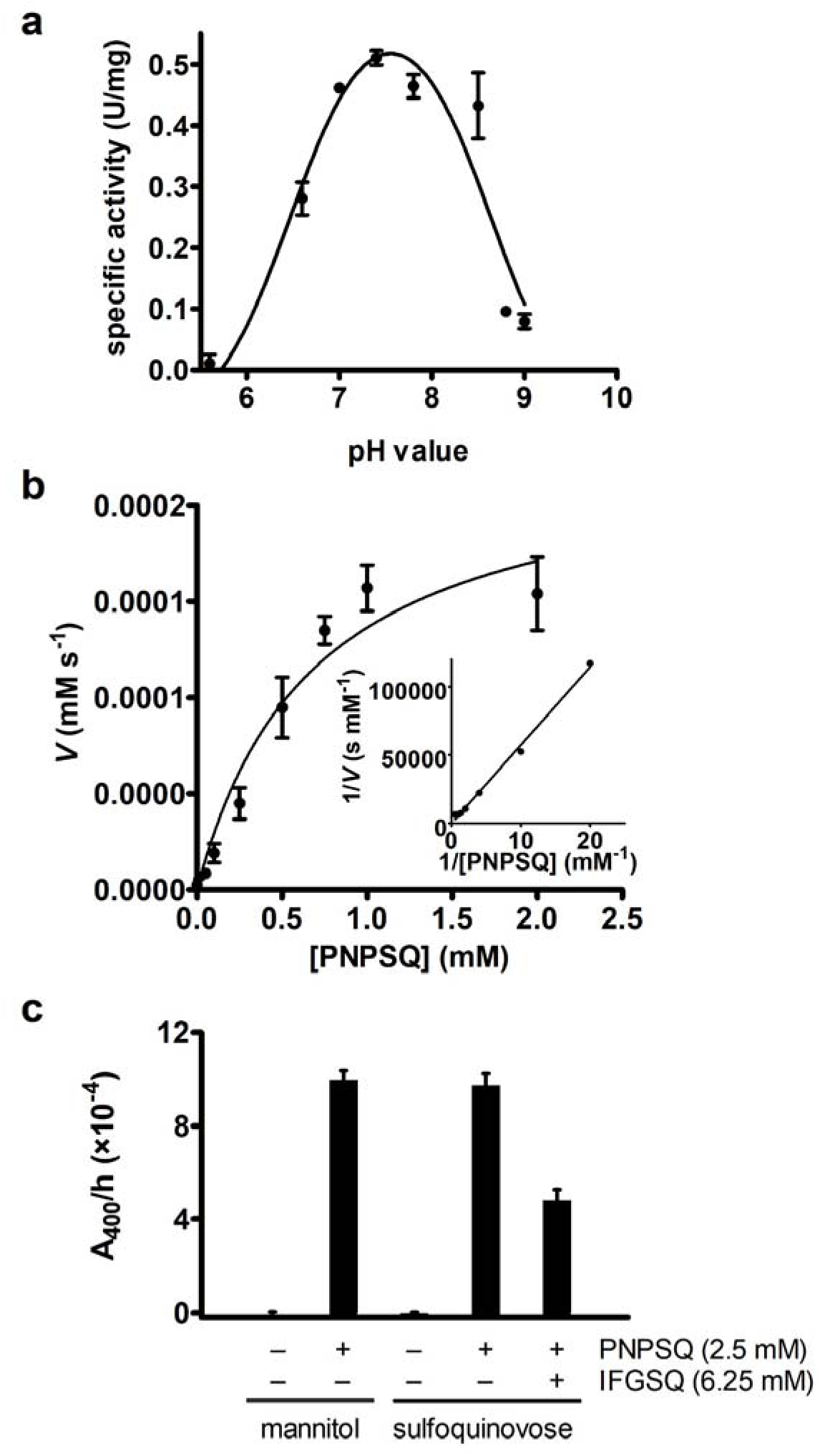
*Rhizobium leguminosarum* SRDI565 produces a functional sulfoquinovosidase that can be detected in cell lysates. a) pH profile of *Rl*SQase*. Specific activities were determined for hydrolysis of PNPSQ. b) Michaelis Menten plot of kinetic parameters for *Rl*SQase* for hydrolysis of PNPSQ. c) Analysis of sulfoquinovosidase activity of *Rl-*SRDI565 lysate grown on sulfoquinovose and mannitol. Cell lysates of soluble proteins derived from growth on SQ or mannitol was standardized for equal protein and SQase activity measured using the chromogenic substrate PNPSQ. SQase activity was confirmed by inhibition by the azasugar inhibitor SGIFG. Error bars denote standard error of the mean.

Direct evidence for enzymatic activity associated with the sulfo-ED operon in *Rl*-SRDI565 was obtained by measuring SQase enzyme activity in cell lysates. The chromogenic substrate 4-nitrophenyl α-sulfoquinovoside (PNPSQ), which was designed as an analogue of the natural substrate SQGro, results in release of the chromophore 4-nitrophenolate, which can be detected using UV-visible spectrophotometry at 400 nm.^23, 24^ *Rl*-SRDI565 was grown to mid-logarithmic phase in Y_5%_M and Y_5%_SQ media, and the harvested cells used to prepare a cell-free lysate containing soluble proteins. Incubation of Y_5%_M and Y_5%_SQ-derived lysates with PNPSQ both resulted in production of 4-nitrophenolate at similar rates. The activity in the YSQ-derived lysate was inhibited by the addition of IFG-SQ, an azasugar inhibitor of SQases that makes key interactions in the active site that mimic those required for substrate recognition (Figure 4c).^24^ The similar levels of activity of SQase in both mannitol and SQ grown *Rl*-SRDI565 is consistent with the abundance of the putative SQase WP_017967311.1 detected by proteomic analysis.

To further confirm that a sulfo-ED pathway was operative in cells, a targeted metabolomics approach was used to detect expected intermediates in bacteria grown on Y_5%_SQ media. Detected intermediates were identified based on their LC-MS/MS retention time and mass spectra with authentic reference standards of the sulfo-EMP and sulfo-ED pathway that were synthesized in-house. Sulfogluconate (SG) was synthesized by oxidation of SQ with iodine^25^ (Figures S11 and S12), while SQ, SF, SFP, DHPS, SLA and SL were prepared as previously reported.^26^ *Rl*-SRDI565 was grown to mid-log phase in Y_5%_M or Y_5%_SQ, metabolically quenched and extracted polar metabolites analyzed by LC/MS-MS. SQ-grown bacteria contained SQ, SF, SG, SL and DHPS, while SFP and SLA could not be detected (Figures 5a-e, Figure S14). The detection of SG is characteristic for a sulfo-ED pathway, and presumably arises from the action of the putative SQ dehydrogenase and SGL lactonase. The identification of DHPS and SF was unexpected, as these intermediates/products of the sulfo-EMP pathway.^11^ BLAST analysis of the genome of *Rl*-SRDI565 did not identify putative genes for the sulfo-EMP pathway. SF may therefore be formed by the action of phosphoglucose isomerase (PGI), while DHPS could be the product of a promiscuous aldehyde reductase. *Rl*-SRDI565 was unable to utilize DHPS or SL as sole carbon source in Y_5%_ medium, supporting the absence of an alternative pathway of sulfoglycolysis that utilizes these intermediates. Unexpectedly, cytosolic levels of DHPS were 20-fold higher than SL, suggesting that cells may lack a membrane transporter to export accumulated DHPS, in contrast to the SL transporter.

**Figure 5:**
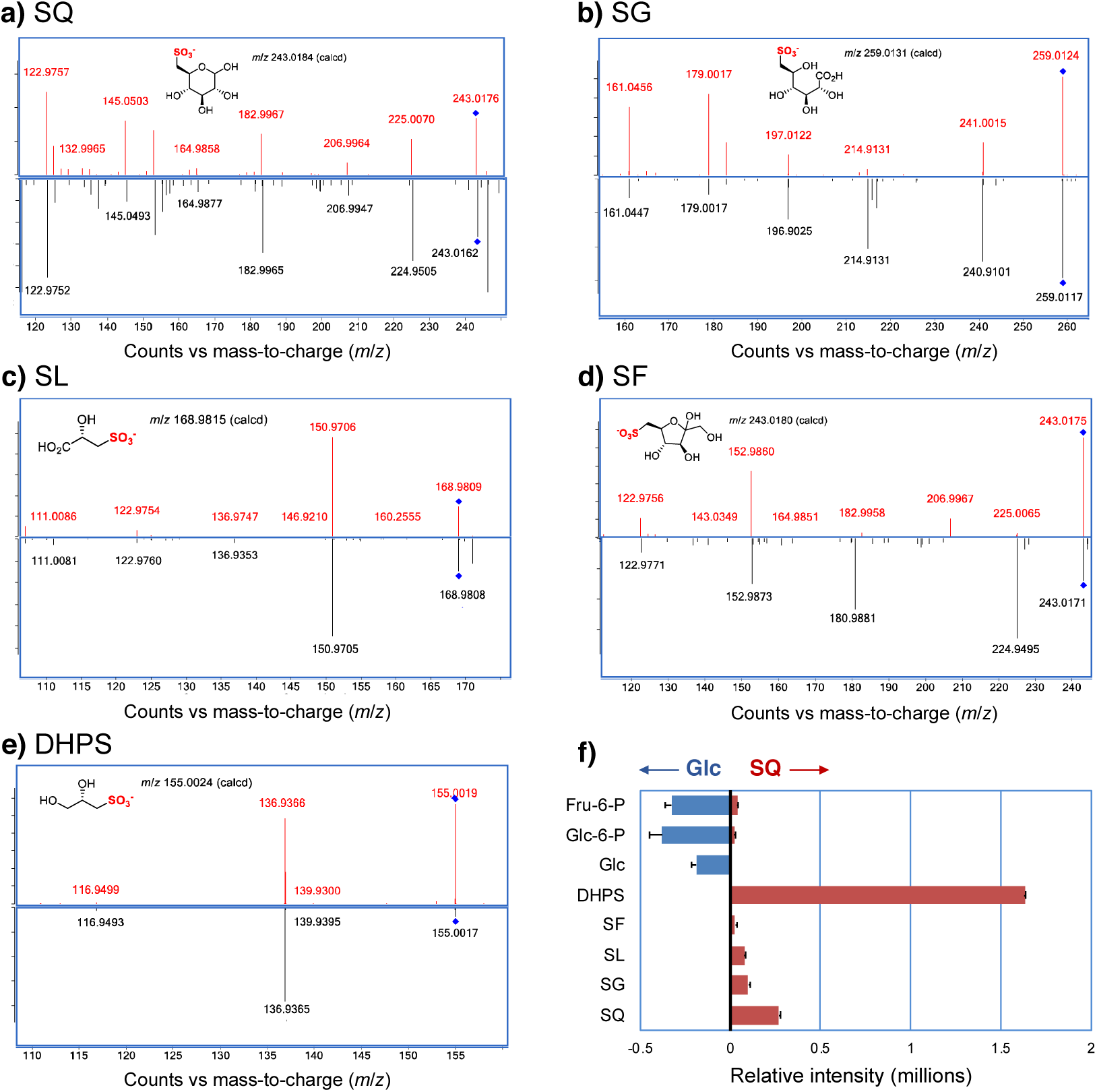
Detection of sulfoglycolytic intermediates and end-products in cytosolic extracts of *Rl*-SRDI565. Rl-SRDI565 was grown on Y_5%_SQ media and metabolically-quenched by rapid cooling to 4 C, followed by extraction of cellular metabolites and lc/ms analysis. Detection of sulfoglycolytic and glycolytic/neoglucogenic intermediates A) SQ, B) SG, C) SL, D) SF, E) DHPS. In each case the upper panel corresponds to the collision-induced dissociation mass spectrum of chemically-synthesized standard, while the lower panel is the equivalent mass spectrum for the metabolite identified in the cytosolic extract. F) Relative mass spectrometric intensities of metabolites from cells grown on Glc or SQ.

NMR and LC-MS/MS analysis of the culture supernatant of both unlabeled and (^13^C_6_)-labelled SQ-cultivated *Rl*-SRDI565 confirmed that the substrate is almost completely consumed by the time bacteria reach stationary growth (final concentration of 0.006±0.001 mM compared to 5.0±0.5 mM SQ in starting medium) (Figure S15). Using a highly sensitive cryoprobe ^13^C NMR spectroscopic analysis revealed that both DHPS and SG were present in culture supernatant of ^13^C_6_-SQ-cultivated *Rl*-SRDI565. Quantitative LC-MS/MS analysis showed that consumption of SQ was associated with production of SL (5.70±0.12 mM), and low levels of DHPS (0.081±0.010 mM), (SG 0.172±0.006 mM) and SF (0.002±0.0001 mM) (Table S2). This experiment was repeated to assess the effect of growth of *Rl*-SRDI565 but using SQGro as carbon source. As noted previously, *Rl*-SRDI565 grows inconsistently on SQGro and complete consumption of SQGro could not be achieved. However, the results of partial consumption broadly agreed with the results for growth on SQ, namely that SL is the major terminal metabolite detected in the culture media, with much lower amounts of SF, SG and DHPS (Table S2).

## Discussion

We demonstrate here that *Rl*-SRDI565 has a functional sulfo-ED pathway that allows these bacteria to utilize SQ as their major carbon source. Catabolism of SQ is primarily or exclusively mediated by a sulfo-ED pathway with production of SL as the major end-product, similar to the situation in *P. putida* SQ1, the only other experimentally described exemplar of this pathway.^12^ In contrast to *P. putida* SQ1, *Rl*-SRDI565 also produces trace amounts of DHPS which could reflect the presence of enzymes which exhibit promiscuous activities similar to those in the conventional sulfo-EMP pathway. This observation is reminiscent of *Klebsiella* sp. strain ABR11 isolated from soil^27^ that is also able to grow on SQ with production of both SL and DHPS. *Klebsiella* sp. strain ABR11 possesses an NAD^+^-specific sulfoquinovose-dehydrogenase activity,^28^ suggesting it has an operative sulfo-ED pathway.

Various bacteria that can metabolize SQ have been isolated from soil including *Agrobacterium* sp.,^28^ *Klebsiella* sp.,^28^ and *Flavobacterium* sp.,^29^ as well as *P. putida* SQ1,^12^ which was isolated from a freshwater littoral sediment. These bacteria may work cooperatively with species such as *Paracoccus pantotrophus* NKNCYSA that can convert SL to mineral sulfur, leading to stoichiometric recovery of sulfite/sulfate.^13^ Together these bacterial communities achieve the complete mineralization of SQ to sulfate, which is available for use by plants.

Proteomic and biochemical evidence suggests that the sulfo-ED pathway is constitutively expressed in *Rl*-SRDI565, and is subject to only limited up-regulation in the presence of SQ. As *Rl*-SRDI565 in the soil is likely to be oligotrophic, constitutive expression of the sulfo-ED pathway may allow simultaneous usage of multiple non-glycolytic substrates without requirement for significant transcriptional changes. Consistent with this view, the proteomic abundance of the putative LacI-type regulator WP_017967302.1 was unchanged between mannitol and SQ grown *Rl*-SRDI565. The sulfo-ED operon in *Rl*-SRDI565 differs from that described for *P. putida* SQ1 through the absence of a putative SQ mutarotase. SQ undergoes mutarotation with a half-life of approximately 6 h, which is much slower than for the glycolytic intermediate Glc-6-P, which has a half-life of just seconds.^20^ Aldose mutarotases are often relatively non-specific and possibly a constitutive mutarotase not in the sulfo-ED operon expressed by the cell provides this catalytic capacity. Alternatively, the SQ dehydrogenase may not be stereospecific, with the ability to act on both anomers of SQ, or even that it acts on α-SQ (the product released from SQGro by an SQase) at a high rate such that mutarotation to β-SQ is of insignificant importance. A second difference in the sulfo-ED operon lies in the presence of an ABC transporter cassette. ABC transporter cassettes are the most common solute transporters, and can translocate their substrates in either a forward or reverse direction.^30^ While we propose that the ABC transporter cassette operates in the forward direction, based on the presence of a signal sequence in the putative solute binding domain targeting it to the periplasm, and consistent with a wide range of sugar import systems, the directionality of transport and thus the choice of substrate (SQ/SQGro versus SL) may depend on the relative abundance of these metabolites intra and extracellularly.

Sulfoglycolysis in *Rl*-SRDI565 leads to production of pyruvate and the excretion of the C3-organosulfonate SL (Figure 6). In order to satisfy the demands of the pentose phosphate pathway and cell wall biogenesis, sulfoglycolytic cells must synthesize glucose-based metabolites such as glucose-6-phosphate and glucose-1-phosphate. Gluconeogenesis has been studied in *Rhizobium leguminosarum* strain MNF3841, and operates through a classical pathway involving fructose bisphosphate aldolase.^31^ Action of phosphoglucose isomerase on SQ might lead to production of SF, thereby explaining the observation of this metabolite in *Rl*-SRDI565. This is not likely to be consequential, as the reversibility of this reaction will ultimately allow complete consumption of any SF through isomerization back to SQ. The formation of DHPS may result from a promiscuous aldehyde reductase. Analysis of spent culture media reveals that the production of DHPS is minor in terms of total carbon balance. However, within the cytosol, DHPS accumulates to levels much higher than SL, presumably because of the absence of a dedicated exporter for the former. Possibly, reduction of SLA to DHPS is reversible and enables conversion of this metabolite to SL and subsequent excretion from the cell. The observation of SG, SF and DHPS in the spent culture media at low levels is suggestive of low levels of leakage of these metabolites from the cell, either through cell lysis or leaky export systems.

**Figure 6:**
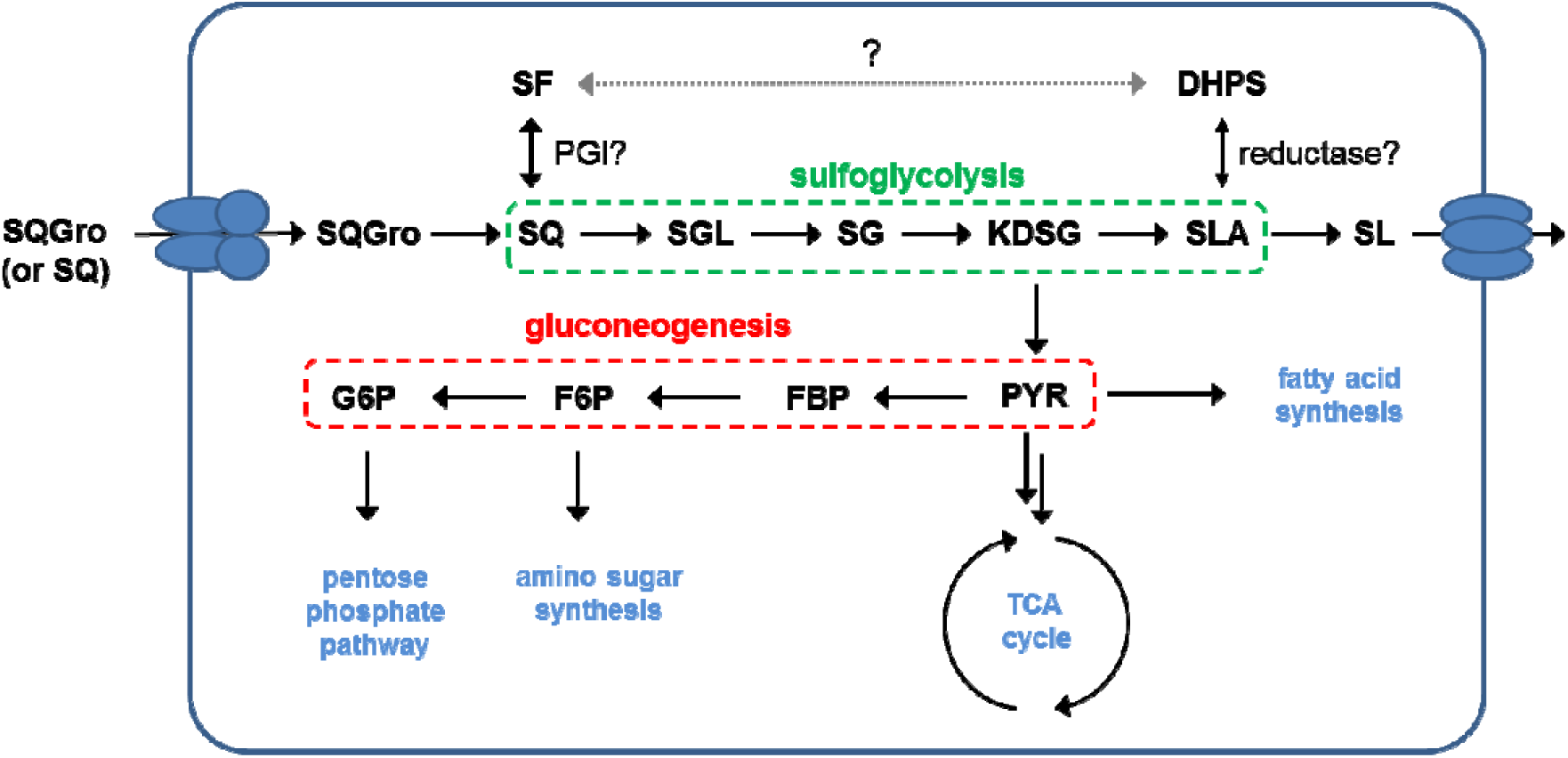
Proposed pathway for SQ metabolism in *Rhizobium leguminosarum* SRDI565.

Given that SQ contains a significant portion of organic sulfur within plants, the pathways of SQ catabolism leading to release of its sulfur may be important to enable recycling of this important macronutrient. Plants can only use sulfate, which is poorly retained by most soils. Biomineralization of organic sulfur to sulfate is important to allow plants to access this element. As one of just two known pathways for the catabolism of SQ, the sulfo-ED pathway is likely to be an important part of environmental breakdown of SQ and may contribute to the persistence of symbiotic rhizobia within the pedosphere. The present work lays the groundwork for a more detailed investigation of sulfoglycolysis in a well-characterized bacterium with an established capability for symbiosis of a leguminous plant host.

## Materials and Methods

### Reagents

SQ, (^13^C_6_)-SQ, SF, SFP, SLA, SL, and DHPS were synthesized as described previously.^26^ IFG-SQ was prepared as described.^24^

### Bacteria and culture conditions

*Rhizobium leguminosarum* bv. *trifolii* SRDI565 was a gift from Dr Ross Ballard, (South Australian Research and Development Institute, Adelaide, South Australia). Minimal salts media consists of 0.5 g·L^−1^ K_2_HPO_4_, 0.2 g·L^−1^ MgSO_4_, 0.1 g·L^−1^ NaCl, 1 M CaCl_2_ 3 mL·L^−1^, adjusted to pH 7.0. YM media consists of minimal salts media plus 1 g·L^−1^ yeast extract, 10 g·L^−1^ mannitol. Y_5%_M consists of minimal salts media plus 50 mg·L^−1^ yeast extract, 5 mM mannitol. Y_5%_SQ consists of minimal salts media plus 50 mg·L^−1^ yeast extract, 5 mM SQ.

Growth curves were determined in a MicrobeMeter built in-house according to published plans^32^ and blueprints available at https://humanetechnologies.co.uk/ The MicrobeMeter was calibrated by performing serial 2-fold dilutions across the detection range of the MicrobeMeter (0-1023 units), starting with an OD_600_ approx. 1 culture of *Rl*-SRDI565. OD_600_ measurements were made with a UV/Vis spectrophotometer and plotted against the reading of the MicrobeMeter. The data was fit to a polynomial to obtain a calibration curve.

#### Proteomic sample preparation

Cells were washed 3 times in PBS and collected by centrifugation at 10,000 × g at 4°C then snap frozen. Frozen whole cell samples were resuspended in 4% SDS, 100 mM Tris pH 8.0, 20 mM DTT and boiled at 95°C with shaking at 2000 rpm for 10 minutes. Samples were then clarified by centrifugation at 17,000 × g for 10 minutes, the supernatant collected, and protein concentration determined by bicinchoninic acid assay (Thermo Scientific Pierce). 100 μg of protein from each sample cleaned up using SP3 based purification according to previous protocols.^33^ Briefly, reduced samples were cooled and then alkylated with 40 mM 2-chloroacetamide (CAA) for 1 hour at RT in the dark. The alkylation reactions were then quenched with 40mM DTT for 10 minutes and then samples precipitated on to SeraMag Speed Beads (GE Healthcare, USA) with ethanol (final concentration 50% v/v). Samples were shaken for 10 minutes to allow complete precipitation onto beads and then washed three times with 80% ethanol. The precipitated protein covered beads were then resuspended in 100mM ammonium bicarbonate containing 2μg trypsin (1/50 w/w) and allowed to digest overnight at 37 °C. Upon completion of the digests samples were spun down at 14000 g for 5 minutes to pellet the beads and the supernatant collected and desalted using homemade C18 stage tips,^34^ then was dried down and stored till analysed by LC-MS.

#### Proteomics analysis using reversed phase LC-MS

Purified peptides prepared were re-suspend in Buffer A* (2% ACN, 0.1% TFA) and separated using a two-column chromatography set up composed of a PepMap100 C18 20 mm × 75 μm trap and a PepMap C18 500 mm × 75 μm analytical column (Thermo Fisher Scientific). Samples were concentrated onto the trap column at 5 μL/min for 5 minutes and infused an Orbitrap Elite™ (Thermo Fisher Scientific). 120 minute gradients were run altering the buffer composition from 1% buffer B (80% ACN, 0.1% FA) to 28% B over 90 minutes, then from 28% B to 40% B over 10 minutes, then from 40% B to 100% B over 2 minutes, the composition was held at 100% B for 3 minutes, and then dropped to 3% B over 5 minutes and held at 3% B for another 10 minutes. The Elite Orbitrap Mass Spectrometers was operated in a data-dependent mode automatically switching between the acquisition of a single Orbitrap MS scan (120,000 resolution) and a maximum of 20 MS-MS scans (CID NCE 35, maximum fill time 100 ms, AGC 1*10^4^).

#### Mass spectrometry data analysis

Proteomic comparison of growth with and without sulfoquinovose was accomplished using MaxQuant (v1.5.5.1).^35^ Searches were performed against *Rhizobium leguminosarum bv. trifolii* SRDI565 (NCBI Taxonomy ID: 935549, downloaded 01-08-2019, 6404 entries) with carbamidomethylation of cysteine set as a fixed modification. Searches were performed with Trypsin cleavage allowing 2 miscleavage events and the variable modifications of oxidation of methionine and acetylation of protein N-termini. The precursor mass tolerance was set to 20 parts-per-million (ppm) for the first search and 10 ppm for the main search, with a maximum false discovery rate (FDR) of 1.0% set for protein and peptide identifications. To enhance the identification of peptides between samples the Match Between Runs option was enabled with a precursor match window set to 2 minutes and an alignment window of 10 minutes. For label-free quantitation, the MaxLFQ option within MaxQuant^36^ was enabled in addition to the re-quantification module. The resulting peptide outputs were processed within the Perseus (v1.4.0.6)^37^ analysis environment to remove reverse matches and common protein contaminates with missing values imputed. The mass spectrometry proteomics data have been deposited to the ProteomeXchange Consortium via the PRIDE partner repository with the dataset identifier PXD015822.

### Chemical synthesis of 6-deoxy-6-sulfo-D-gluconate (SG)

NaOH in methanol (4% w/v, 4 mL) was added dropwise to a stirred solution of sulfoquinovose (100 mg, 0.410 mmol) and iodine (209 mg, 1.65 mmol) in water (1 mL) and methanol (4 mL) held at 40 °C. As the sodium hydroxide was added the color of iodine dissipated. The solvent was evaporated under reduced pressure and the crude residue was subjected to flash chromatography (EtOAc/MeOH/H_2_O, 4:2:1 to 2:2:1, then water) to give the 6-deoxy-6-sulfogluconate sodium salt (89.2 mg). An aqueous solution of the sodium salt was eluted through a column of Amberlite IR120 (H^+^ form) resin. The acidic eluate was collected and concentrated under reduced pressure give SG (71.3 mg, 67%). ^1^H NMR (400 MHz, D_2_O): δ 4.23–4.15 (1 H, m, H2), 4.13 (1 H, d, *J* = 3.3 Hz, H3), 4.05 (1 H, t, *J* = 3.2 Hz, H5), 3.74 (1 H, dd, *J* = 6.5, 3.4 Hz, H4), 3.35 (1 H, d, *J* = 14.6 Hz, H6a), 3.05 (1 H, dd, *J* = 14.6, 9.7 Hz, H6b); ^13^C{^1^H} NMR (100 MHz, D_2_O) δ 178.7 (C1), 74.2 (C4), 73.8 (C2), 70.8 (C3), 67.8 (C5), 53.4 (C6); HRMS (ESI^−^) calcd for C_6_H_11_O_9_S [M^−^] 259.0129, found 259.0131.

### Metabolite analysis of *Rhizobium leguminosarum* cell extracts

#### Metabolic quenching and extraction

*Rl*-SRDI565 was grown on Y_5%_SQ or Y_5%_ containing 35 mM glucose to mid-logarithmic phase (approx. 0.15), as calculated based on the OD_600_ measured by Cary 50 UV/visible spectrophotometer, and were rapidly quenched in a prechilled 15 mL Falcon tube containing phosphate buffered saline (PBS) at 4 °C. Ice-cold PBS (11 mL) was infused into cell culture media (4 mL). The Falcon tubes were mixed by inversion and incubated in ice/water slurry for 5 min then were centrifuged at 2000 × *g* at 1 °C for 10 min. The supernatant was removed by aspiration and cell pellets were washed twice with 1 mL of ice-cold PBS (with resuspension each time) and transferred into 1.5 mL Eppendorf tubes. Cells were pelleted by centrifugation at 14000 rpm and residual solvent was carefully removed. Cell pellets were stored at -80°C until extraction. Cells were extracted in 200 µL of extraction solution (methanol/water, 3:1 v/v) containing an internal standard, 5 µM ^13^C_4_-aspartate (Cambridge Isotopes), and subjected to 10 freeze-thaw cycles to facilitate cell lysis (30 s in liquid nitrogen, followed by 30 s in dry ice/ethanol bath). Debris was pelleted by centrifugation at 14000 rpm, 5 min, 1°C and cell lysate was transferred into a HPLC vial insert for LC/MS analysis.

#### LC/MS analysis and identification of sulfonate metabolites

Separation and detection of polar metabolites was performed using an Agilent Technologies 1200 series high performance liquid chromatography (HPLC) coupled to a quadrupole time-of-flight mass spectrometer (6545 QTOF, Agilent Technologies) using a method modified from Masukagami *et al*.^38^ Metabolite extracts were transferred into 2 mL auto sampler vials with glass inserts and placed in the auto sampler kept at 4 °C prior to analysis. Metabolite separation was performed by injecting 7 µL of the extract into a SeQuant® ZIC-pHILIC PEEK coated column (150 mm × 4.6 mm, 5 µm polymer, Merck Millipore) maintained at 25°C, with a gradient of solvent A (20 mM ammonium carbonate, pH 9.0, Sigma-Aldrich) and solvent B (100% acetonitrile, Hypergrade for LCMS LiChrosolv, Merck) at a flow rate of 0.3 mL/min. A 33.0 min gradient was setup with time (*t*) = 0 min, 80% B; *t* = 0.5 min, 80% B; *t* = 15.5 min, 50% B; *t* = 17.5 min, 30% B; *t* = 18.5 min, 5% B; *t* = 21.0 min, 5% B; *t* = 23.0 min, 80% B.

The LC flow was directed into an electrospray ionization (ESI) source with a capillary voltage of 2500 V operating in negative ionization mode. Drying nitrogen gas flow was set to 10 L/min, sheath gas temperature and nebulizer pressure were set to 300 °C and 20 psi, respectively. The voltages of fragmentor and skimmer were set at 125 V and 45 V, respectively. Data was acquired in MS and MS/MS mode, with a scan range of 60 to 1700 *m/z* and 100 to 1700 *m/z* respectively, at a rate of 1.5 spectra/sec. MS/MS acquisition was performed with four collision energies (0, 10, 20 and 40 V). The mass spectrometer was calibrated in negative mode prior to data acquisition and mass accuracy during runs was ensured by a continuous infusion of reference mass solution at a flow rate of 0.06 mL/min (API-TOF Reference Mass Solution Kit, Agilent Technologies). Data quality was ensured by multiple injections of standards (with 1.5 µM concentration each) and pooled biological sample (a composite of cell extracts) used to monitor the instrument performance. Samples were randomized prior to metabolite extraction and LC/MS analysis.

#### Standard preparation

Standards of selected metabolites (Supplementary Table 1) were prepared at 10 µM in 80% acetonitrile (Hypergrade for LCMS LiChrosolv, Merck) and injected separately into a column connected to mass spectrometer interface. Retention time and detected molecular ion were used to create a targeted MS/MS acquisition method. The spectra, mass to charge (*m/z*) and retention time (RT) were imported into a personal compound database and library (PCDL Manager, version B.07.00, Agilent Technologies) used in data processing workflow.

#### Data analysis

Data were analysed using MassHunter Qualitative and Quantitative Analysis software (version B.07.00, Agilent Technologies). Identification of metabolites was performed in accordance with metabolite identification (Metabolomics Standard Initiative, MSI) level 1 based on the retention time and molecular masses matching to authentic standards included in the personal database and library. Peak integration was performed in MassHunter quantitative software (version B.07.00, Agilent Technologies) on the spectra from identified metabolites.

#### Cloning, expression and kinetic analysis of *Rl-*SRDI565 sulfoquinovosidase (*Rl*SQase*)

The gene sequence coding for *Rl*SQase* SERp mutant was synthesised with codon optimisation for expression in *E. coli* and was cloned within a pET-28a(+) vector with C-terminal His-tag through GenScript. The plasmid His_6_-*Rl*SQase*-pET-28a(+) containing the gene for target *Rl*SQase* was transformed into *E. coli* BL21(DE3) cells for protein expression. Pre-cultures were grown in LB-medium (5 mL) containing 30 µg/mL for 18 h at 37 °C, 200 rpm. Cultures (1 L LB-medium supplemented with kanamycin 30 µg/mL) were inoculated with the pre-culture (5 mL) and incubated at 37 °C, 200 rpm until an OD_600_ of 0.6- 0.8 was achieved. Protein expression was induced by addition of IPTG (1 mM) and shaking was continued overnight (20-22 h) at 18 °C, 200 rpm. The cells were harvested by centrifugation (5000 rpm, 4 °C, 20 min), resuspended in 50 mM Tris, 300 mM NaCl pH 7.5 buffer and were subjected to further cell lysis. Cells were disrupted using French press under 20 k Psi pressure and the lysate was centrifuged at 50,000 g for 30 min.

The N-terminal His_6_-tagged protein was purified by immobilized metal ion affinity chromatography, followed by size exclusion chromatography (SEC) (Figure S10a). The lysate was loaded onto a pre-equilibrated Ni-NTA column, followed by washing with load buffer (50 mM Tris-HCl, 300 mM NaCl, 30 mM imidazole pH 7.5). The bound protein was eluted using a linear gradient with buffer containing 500 mM imidazole. Protein containing fractions were pooled, concentrated and loaded onto a HiLoad 16/600 Superdex 200 gel filtration column pre-equilibrated with 50 mM Tris-HCl, 300 mM NaCl pH 7.5 buffer. The protein was concentrated to a final concentration of 60 mg mL^−1^ using a Vivaspin® 6 with a 300 kDa MW cut-off membrane for characterization and enzyme assays.

### SEC-MALS analysis

Experiments were conducted on a system comprising a Wyatt HELEOS-II multi-angle light scattering detector and a Wyatt rEX refractive index detector linked to a Shimadzu HPLC system (SPD-20A UV detector, LC20-AD isocratic pump system, DGU-20A3 degasser and SIL-20A autosampler). Experiments were conducted at room temperature (20 ± 2°C). Solvents were filtered through a 0.2 µm filter prior to use and a 0.1 µm filter was present in the flow path. The column was equilibrated with at least 2 column volumes of buffer (50 mM Tris, 300 mM NaCl pH 7.5) before use and buffer was infused at the working flow rate until baselines for UV, light scattering and refractive index detectors were all stable. The sample injection volume was 100 µL *Rl*SQase* at 6 mg/mL in 50 mM Tris buffer, 300 mM NaCl pH 7.5. Shimadzu LC Solutions software was used to control the HPLC and Astra V software for the HELEOS-II and rEX detectors (Figure S10b). The Astra data collection was 1 min shorter than the LC solutions run to maintain synchronisation. Blank buffer injections were used as appropriate to check for carry-over between sample runs. Data were analysed using the Astra V software. Molar masses were estimated using the Zimm fit method with degree 1. A value of 0.158 was used for protein refractive index increment (dn/dc).

### Enzyme kinetics of *Rl*SQase

#### Michaelis Menten plot

Kinetic analysis of *Rl*SQase* was performed using PNPSQ as substrate, using a UV/visible spectrophotometer to measure the release of the 4-nitrophenolate (λ = 348 nm). Assays were carried out in 50 mM sodium phosphate, 150 mM NaCl, pH 7.2 at 30 °C using 212 nM *Rl*SQase* at substrate concentrations ranging from 0.05 μM to 4 mM. Using the extinction coefficient for 4-nitrophenolate of 5.125 mM^−1^ cm^−1^, kinetic parameters were calculated using Prism.

#### pH profile

For the determination of pH profile, specific activities of *Rl*SQase* were monitored by measuring absorbance changes at λ = 348 nm in the presence of the following buffers: sodium acetate buffer (pH 5.6, sodium phosphate buffer (pH 6.0–8.5), and glycine NaOH buffer (pH 8.8–9.2). The assays were performed at 30 °C in duplicates and specific activities determined using extinction coefficient of PNP at isobestic point (348 nM) as 5.125 mM^−1^ cm^−1^ (Supplementary Figure Sx). One unit of SQase activity is defined as the amount of protein that releases 1 μmol PNP per minute.

### Detection of SQase activity in cell lysates

*Rl-*SRDI565 was grown in 50 mL Y_5%_M and Y_5%_SQ media at 30 □ to mid log phase, approximately OD_600_ = 0.2, measured using a Varian Cary50 UV/visible spectrophotometer. Cells were harvested by adding 3× volume of ice-cold PBS to metabolically quench the samples then centrifuged at 2000 *g*, 4 °C for 10 min. The supernatant was discarded and the cells were washed 3 times with ice-cold PBS, with each wash involving resuspension and centrifugation at 2000 *g*, 4 °C for 10 min. The cells were collected once more by centrifugation at 10,000 *g*, 4 °C, for 1 min then snap frozen in liquid nitrogen and stored at - 80 °C.

Cells were lysed by addition of 1000 μL pre-chilled PBS, 1 μL RNaseA, 1 μL DNase, 1 μL 100 mg·mL^−1^ hen egg white lysozyme (Sigma), and a 1× final concentration of cOmplete EDTA-free protease inhibitor cocktail (Roche) to the cell pellet. The cells were gently resuspended and mixed at 4 °C for 10 min. The suspension was placed on ice and irradiated with a Sonoplus HD3200 MS 73 sonicator probe (Bandelin) at a frequency of 20 kHz, 20% amplitude, pulse 2s on 8s off, repeated for a total time of the sonication to 150 s, then incubated on ice for 5 min. The suspension was clarified by centrifuging at 14,000 *g*, 4 □ for 1 min and the supernatant was filtered through a Nanosep mini centrifugal spin column with a 0.2 μm filter (Pall) into a 1.5mL Eppendorf tubes and stored at 4 °C. Protein concentration was determined using a BCA assay.

SQase activity was measured in triplicate using PNPSQ and an Agilent Cary UV Workstation (G5191-64000) at 30□. Reactions contained buffer consisted of 50 mM NaPi and 150 mM NaCl, pH=7.4, and 2.5 mM PNPSQ. Reactions were initiated by addition of SQ- or mannitol-derived lysate to a final concentration of 43.7 μg·mL^−1^ protein, and absorbance was monitored at 400 nm for 3 h. After 3 h, IFGSQ was added to a final concentration of 6.25 mM to the SQ-lysate sample, and absorption monitored for 3 h.

### Quantitation of metabolite levels in spent culture media

The metabolites (DHPS, SF, SQ, SL and SG) present in spent culture media were quantified against standard solutions of pure metabolites by HPLC-ESI-MS/MS. Quantification was done with the aid of calibration curves generated by dissolving the pure standards in spent media from *Rl*-SRDI565 grown on Y_5%_M. Spiked spent media was diluted 100-fold with water and then analysed by LC-MS/MS with α-MeSQ as internal standard. For experimental determination of metabolites, spent culture media from *Rl*-SRDI565 grown in Y_5%_SQ or Y_5%_SQGro were diluted 100-fold with water and analysed by LC-MS/MS with α-MeSQ as internal standard.

HPLC-ESI-MS/MS analysis was performed using a TSQ Altis triple quadrupole mass spectrometer (Thermo Fisher Scientific) coupled with a Vanquish Horizon UHPLC system (Thermo Fisher Scientific). The column was a ZIC-HILIC column (5 µm, 50 × 2.1 mm; Merck). The HPLC conditions were: from 90% B to 40% B over 15 min; then 40% B for 5 min; back to 90% B over 1 min (solvent A: 20 mM NH_4_OAc in 1% acetonitrile; solvent B: acetonitrile); flow rate, 0.30 ml min^−1^; injection volume, 1 µl. The mass spectrometer was operated in negative ionization mode. Quantitation was done using the MS/MS selected reaction monitoring (SRM) mode using Thermo Scientific XCalibur software and normalized with respect to the internal standard, α-MeSQ. Prior to analysis, for each analyte, the sensitivity for each SRM-MS/MS transition was optimized.

DHPS: ESI–MS/MS *m*/*z* of [M-H]^-^ 155, product ions 137, 95; retention time: 4.91 min α-MeSQ (internal standard): ESI–MS/MS *m*/*z* of [M-H]^-^ 257, product ions 166, 81; retention time: 6.31 min

SF: ESI–MS/MS *m*/*z* of [M-H]^-^ 243, product ions 207, 153; retention time: 6.81 min

SQ: ESI–MS/MS *m*/*z* of [M-H]^-^ 243, product ions 183, 123; retention time: 7.58 and 7.89 min for α / β

SL: ESI–MS/MS *m*/*z* of [M-H]^-^ 169, product ions 107, 71; retention time: 9.26 min

SG: ESI–MS/MS *m*/*z* of [M-H]^-^ 259, product ions 241, 161; retention time: 9.66 min

SQGro: ESI–MS/MS *m*/*z* of [M-H]^-^ 317, product ions 225, 165; retention time: 7.15 min

## Acknowledgements

This work was supported by grants from the Australian Research Council (DP180101957), the National Health and Medical Research Council of Australia (APP1100164, GNT1139549) and the Leverhulme Trust; support from The Walter and Eliza Hall Institute of Medical Research, the Australian Cancer Research Fund, and a Victorian State Government Operational Infrastructure support grant. MJM is an NHMRC Principal Research Fellow, G.J.D. is a Royal Society Ken Murray Research Fellow. JL is supported by a PhD scholarship from the Chinese Scholarship Council. We thank Humane Technologies for support with the MicrobeMeter, the Melbourne Mass Spectrometry and Proteomics Facility of the Bio21 Institute at the University of Melbourne, Palika Abayakoon and Janice Mui for reagents, and Dr Shuai Nie, Yunyang Zhang and Alex Chen (Thermo Fisher) for technical support. Thermo Fisher Scientific Australia are acknowledged for access to the TSQ Altis triple quadrupole mass spectrometer.

