## Supplementary information for "A sulfoglycolytic Entner-Doudoroff pathway in *Rhizobium leguminosarum* bv. *trifolii* SRDI565"

**Table S1.** LC-MS data for glycolytic and sulfoglycolytic standards.

| name | acronym | formula | *m/z* | retention time |
|---|---|---|---|---|
| sulfoquinovose | SQ | $C_6H_{12}O_8S$ | 243.01801 | 19.54 |
| 6-deoxy-6-sulfogluconate | SG | $C_6H_{12}O_9S$ | 259.0129 | 20.91 |
| 3-sulfolactaldehyde | SLA | $C_3H_6O_5S$ | 152.9863 | 17.647 |
| 3-sulfolactate | SL | $C_3H_6O_6S$ | 168.9812 | 20.379 |
| 2,3-dihydroxypropanesulfonate | DHPS | $C_3H_8O_5S$ | 155.002 | 14.457 |
| 6-deoxy-6-sulfofructose | SF | $C_6H_{12}O_8S$ | 243.018 | 17.623 |
| glucose | Glc | $C_6H_{12}O_6$ | 179.0561 | 16.58 |
| glucose-6-phosphate | Glc-6-P | $C_6H_{13}O_9P$ | 259.0224 | 19.9 |
| fructose 6-phosphate | Fru-6-P | $C_6H_{13}O_9P$ | 259.0224 | 18.897 |

**Table S2.** Analysis of sulfonate metabolites detected in spent culture media of *Rl*-SRDI565 grown on 5.0±0.5 mM SQ or SQGro (standard error estimate). Measurements were performed in triplicate using LC/MS-MS. Errors listed in the table are standard error mean.

| | metabolite concentration (mM) | |
|---|---|---|
| **metabolite** | **spent media (SQ)** | **spent media (SQGro)**[a] |
| SL | 5.696 ± 0.119 | 3.143 ± 0.033 |
| DHPS | 0.081 ± 0.010 | 0.116 ± 0.002 |
| SQ | 0.006 ± 0.001 | 0.215 ± 0.001 |
| SF | 0.002 ± 0.0001 | 0.003 ± 0.0001 |
| SG | 0.172 ± 0.006 | 0.200 ± 0.008 |
| SQGro | – | 2.165 ± 0.056 |

[a] Growth on SQGro was incomplete.

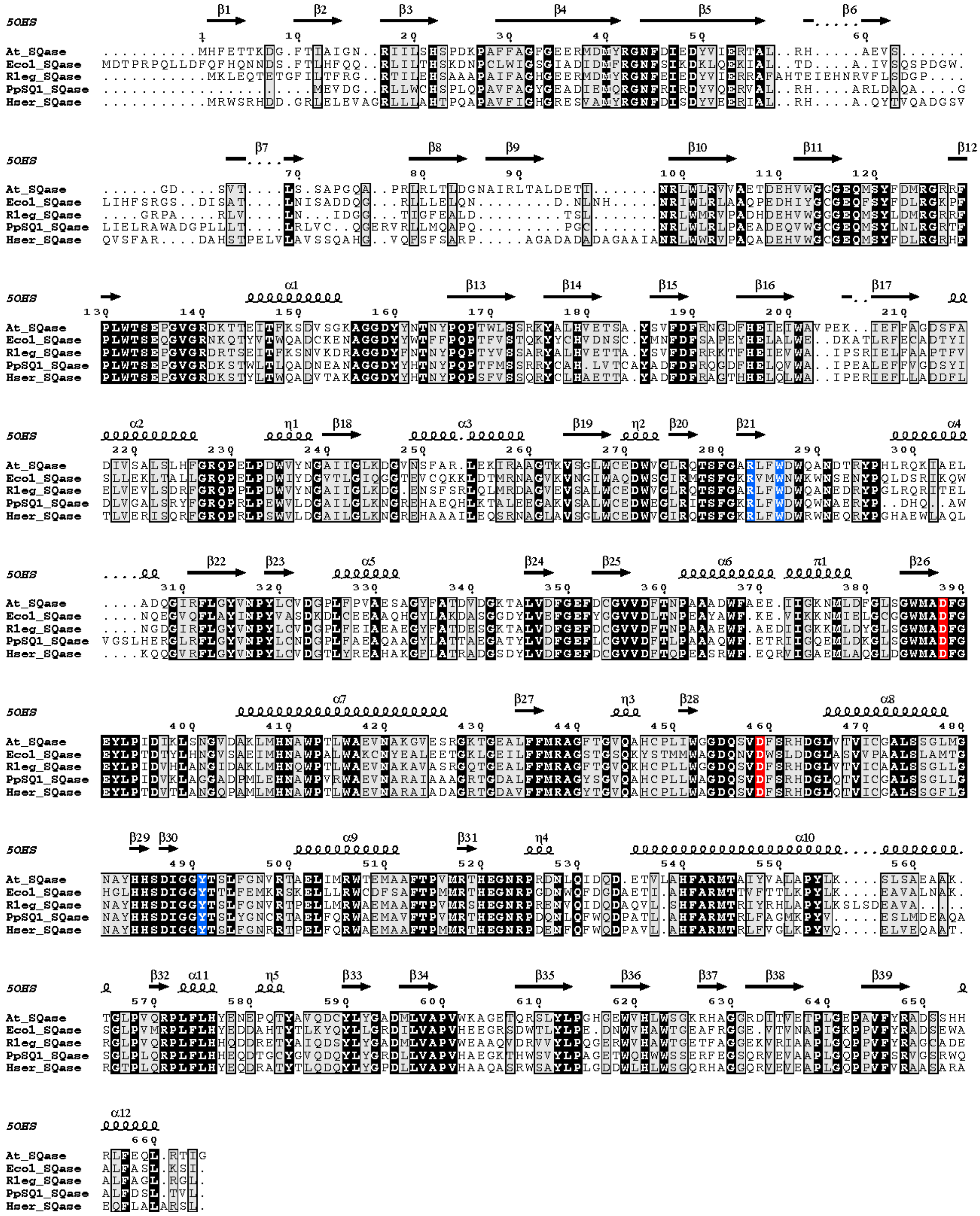

**Figure S1.** Alignment of various putative sulfoquinovosidases (SQase) showing secondary structural elements. Sequences are: At_SQase, Atu3285 (sulfoquinovosidase, WP_010972911.1) from *Agrobacterium tumefaciens* str. C58; Rleg_SQase; Ecol_SQase, YihQ (sulfoquinovosidase, NP_418314.1) from *Esherichia coli*; RLEG16_RS0123470 (putative sulfoquinovosidase, WP_017967311.1) from *Rhizobium leguminosarum* SRDI565; PpSQ1_SQase, PpSQ1_RS00420 (putative sulfoquinovosidase, WP_052326100.1) from *Pseudomonas putida* SQ1; Hser_SQase, hsc_RS17980 (putative sulfoquinovosidase, WP_069374724.1) from *Herbaspirillum seropedicae* AU14040. The secondary structural elements are annotated from the structure of At_SQase, (PDB 5OHS). Shown in red are conserved residues shown to be important in the catalytic mechanism. Shown in blue are residues that have been implicated in binding the sulfonate of the substrate.(1)

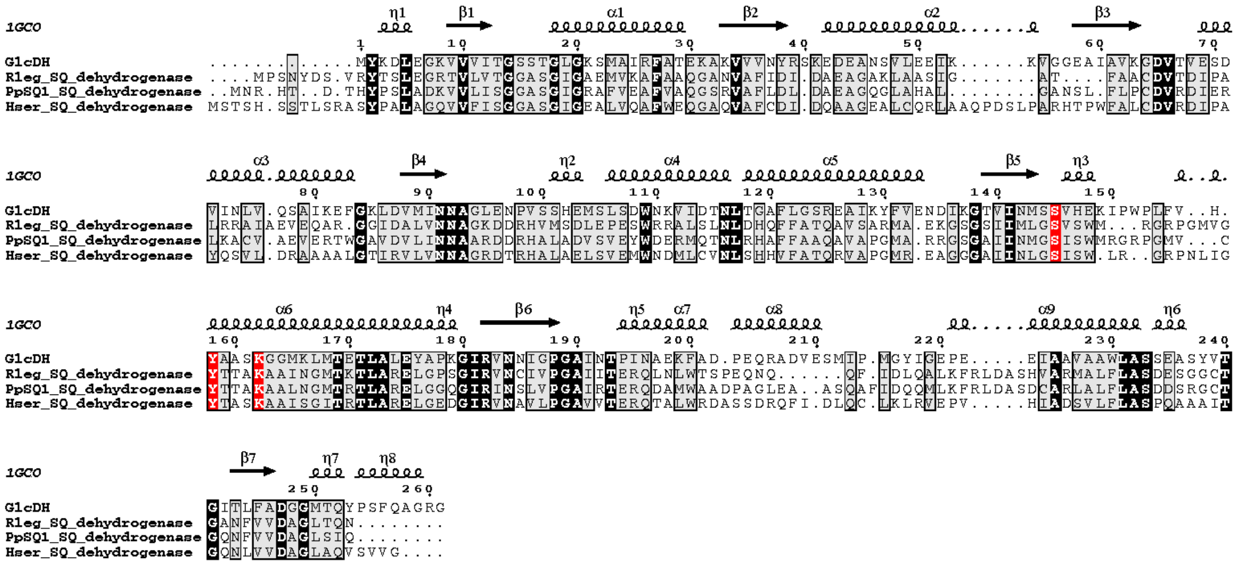


**Figure S2.** Alignment of various putative NAD$^+$-dependent sulfoquinovose dehydrogenases (SQ dehydrogenase) showing secondary structural elements. Sequences are: GlcDH (glucose dehydrogense, BAA01475.1) from *Bacillus megaterium*; Rleg_SQ_dehydrogenase, RLEG16_RS0123465 (putative SQ dehydrogenase, WP_017967310.1) from *Rhizobium leguminosarum* SRDI565; PpSQ1_SQ_dehydrogenase, PpSQ1_RS00400 (putative SQ dehydrogenase, WP_039601086.1) from *Pseudomonas putida* SQ1; Hser_SQ_dehydrogenase, hsc_RS17955 (putative SQ dehydrogenase, WP_069374720.1) from *Herbaspirillum seropedicae* AU14040. The secondary structural elements are annotated from the structure of GlcDH (PDB 1GCO). Shown in red are conserved residues shown to be important in the catalytic mechanism.(2)

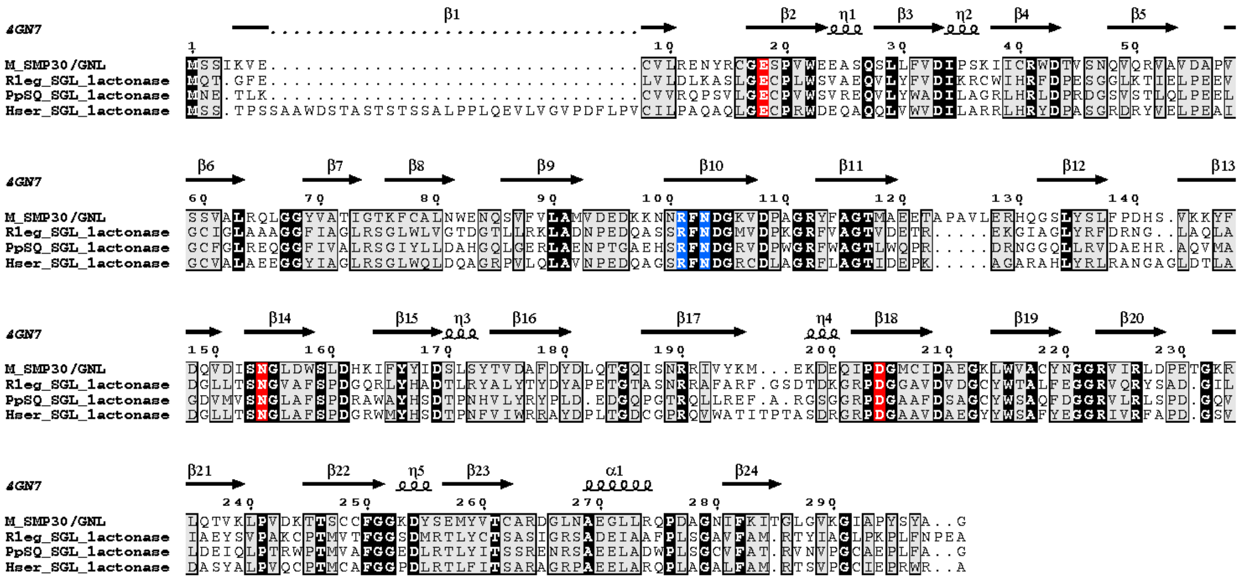

**Figure S3.** Alignment of various putative 6-deoxy-6-sulfogluconolactone lactonases (SGL lactonase) showing secondary structural elements. Sequences are:, M_SMP30/GNL (mouse senescence marker protein-30, gluconolactonase) from mouse; Rleg_SGL_lactonase, RLEG16_RS0123460 (putative SGL lactonase, WP_017967309.1) from *Rhizobium leguminosarum* SRDI565; PpSQ1_SGL_lactonase, PpSQ1_RS00405 (putative SGL lactonase, WP_039601087.1) from *Pseudomonas putida* SQ1; Hser_SGL_lactonase, hsc_RS17995 (putative SGL lactonase, WP_083247167.1) from *Herbaspirillum seropedicae* AU14040. The secondary structural elements are annotated from the structure of M_SMP30/GNL (PDB 4GN7). Shown in red are conserved residues shown to be important in the catalytic mechanism. Shown in blue are residues that have been implicated in substrate binding.(3)

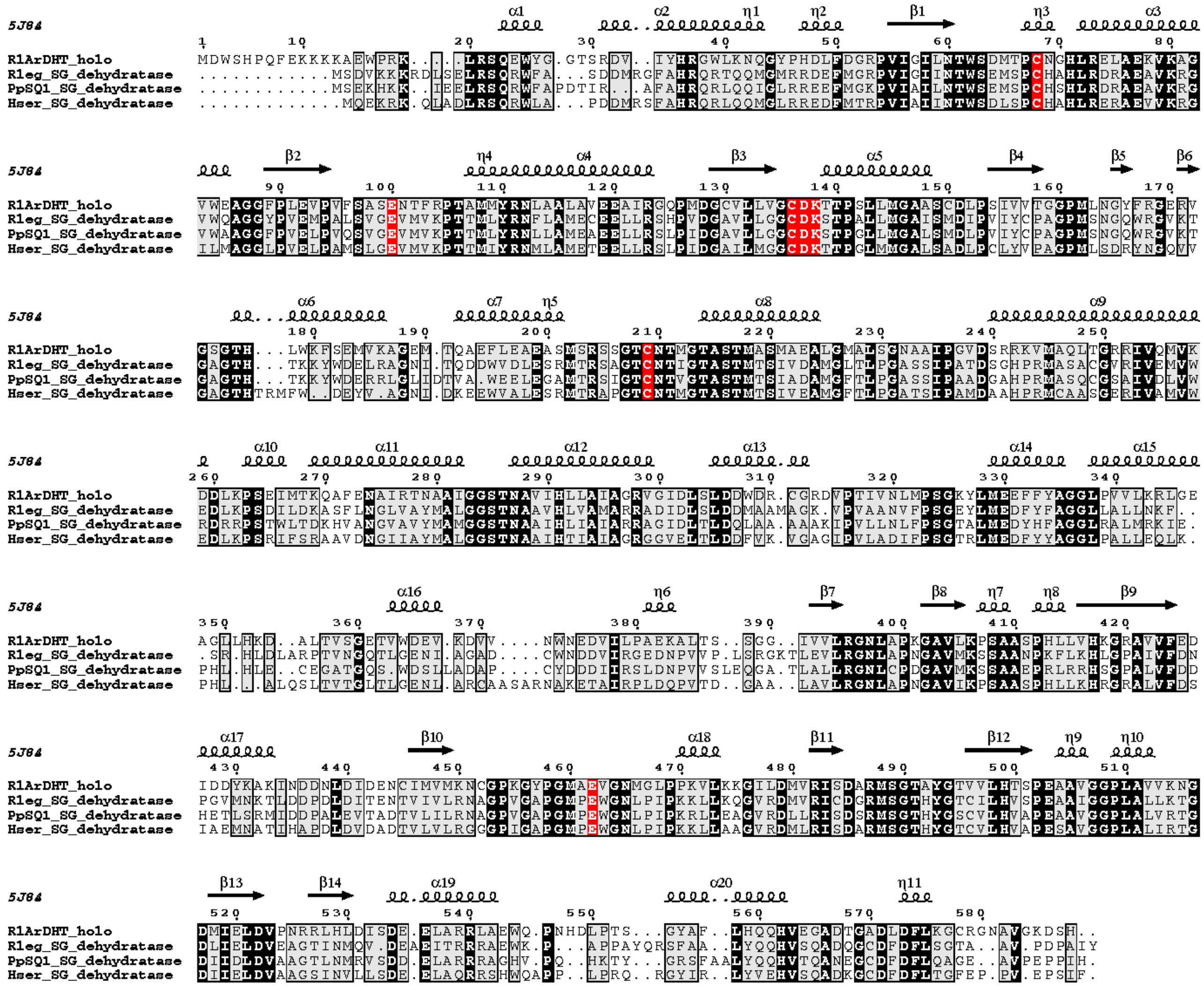


**Figure S4.** Alignment of various putative 6-deoxy-6-sulfogluconate dehydratases (SG dehydratase) showing secondary structural elements. Sequences are: RlArDHT_holo, Rleg2_2909 (L-arabonate dehydratase, ANS60459.1) from *Rhizobium leguminosarum bv. trifolii* WSM2304; Rleg_SG_dehydratase, RLEG16_RS0123450 (putative SG dehydratase, WP_017967307.1) from *Rhizobium leguminosarum* SRDI565; PpSQ1_SG_dehydratase, PpSQ1_RS00395 (putative SG dehydratase,WP_039601085.1) from *Pseudomonas putida* SQ1; Hser_SG_dehydratase, hsc_RS17970 (putative SG dehydratase, WP_069374722.1) from *Herbaspirillum seropedicae* AU14040. The secondary structural elements are annotated from the structure of RlArDHT_holo (PDB 5J84). Shown in red are conserved residues shown to be important in the catalytic mechanism, for formation of a [2Fe-2S] cluster, or for coordination with $Mg^{2+}$.(4)

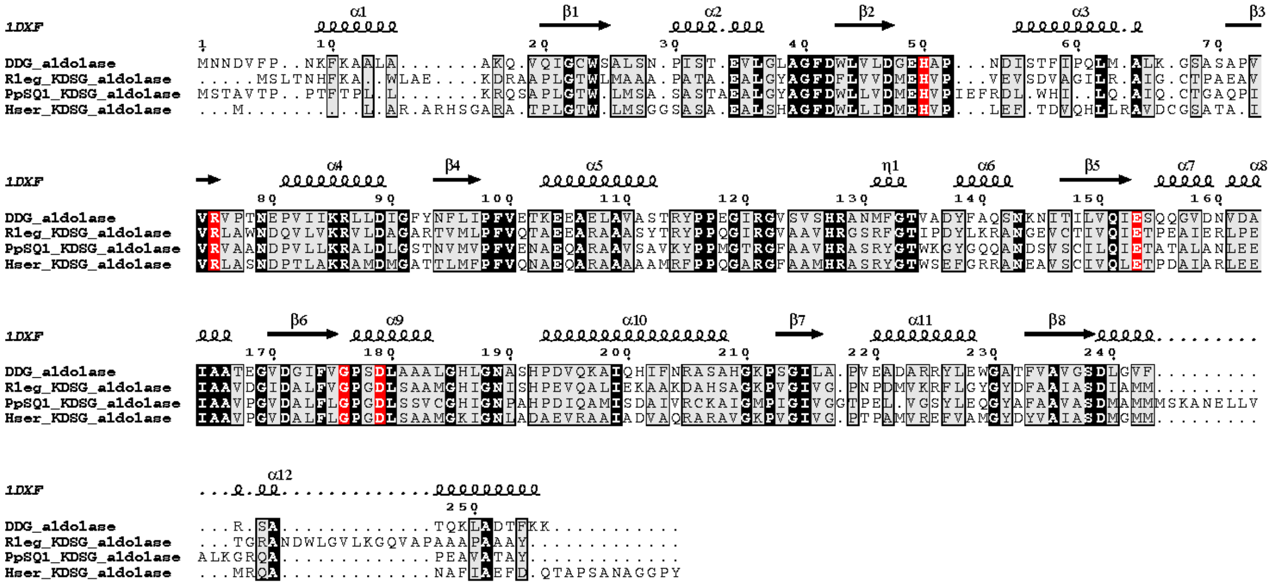


**Figure S5.** Alignment of various putative 2-keto-3,6-dideoxy-6-sulfogluconate aldolases (putative KDSG aldolase) showing secondary structural elements. Sequences are: DDG_ aldolase, ECET_16830 (2-dehydro-3-deoxy-galactarate aldolase, WP_001058209.1) from *Escherichia coli*; Rleg_KDSG_aldolase, RLEG16_RS0123455 (putative KDSG aldolase, WP_017967308.1) from *Rhizobium leguminosarum* SRDI565; PpSQ1_KDSG_aldolase, PpSQ1_RS00450 (putative KDSG aldolase, WP_052326102.1) from *Pseudomonas putida* SQ1; Hser_KDSG_aldolase, hsc_RS17965 (putative KDSG aldolase, WP_069375604.1) from *Herbaspirillum seropedicae* AU14040. The secondary structural elements are annotated from the structure of DDG aldolase (PDB 1DXF). Shown in red are conserved residues shown to be important in the catalytic mechanism or binding a divalent metal centre.(5)

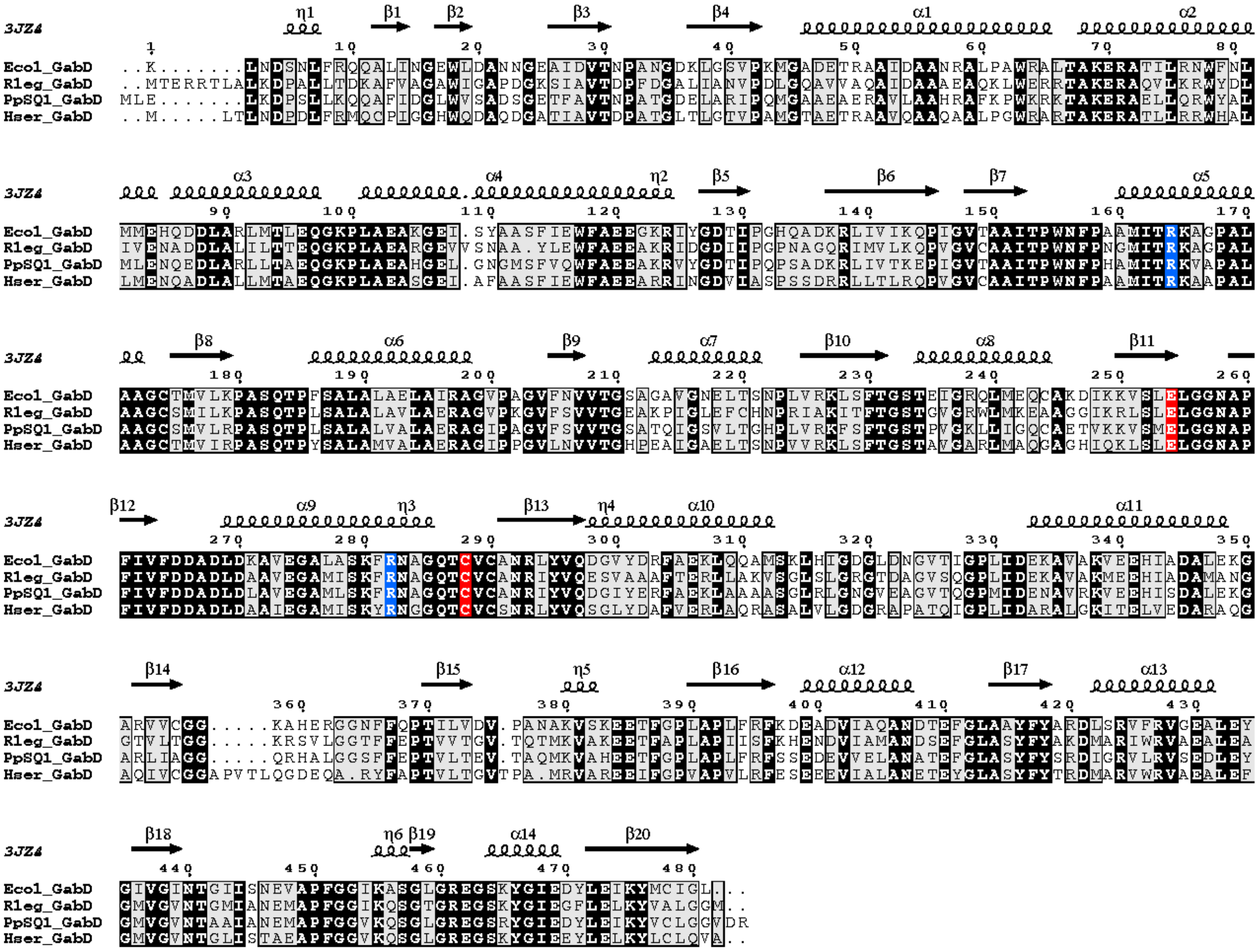


**Figure S6.** Alignment of various putative sulfolactaldehyde dehydrogenases (GabD) showing secondary structural elements. Sequences are: Ecol_GabD, ECs3522 (succinate semialdehyde reductase, NP_311549.1) from *Escherichia coli*; Rleg_GabD, GabD (putative SLA dehydrogenase, WP_017967313.1) from *Rhizobium leguminosarum* SRDI565; PpSQ1_GabD, PpSQ1_RS00390 (putative SLA dehydrogenase, WP_039601084.1) from *Pseudomonas putida* SQ1; Hser_GabD, GabD (putative SLA dehydrogenase, WP_069374726.1) from *Herbaspirillum seropedicae* AU14040. The secondary structural elements are annotated from the structure of Ecol_GabD (PDB 3JZ4). Shown in red are conserved residues shown to be important in the catalytic mechanism. Shown in blue are residues that have been implicated in binding the sulfonate of the substrate.(6,7)

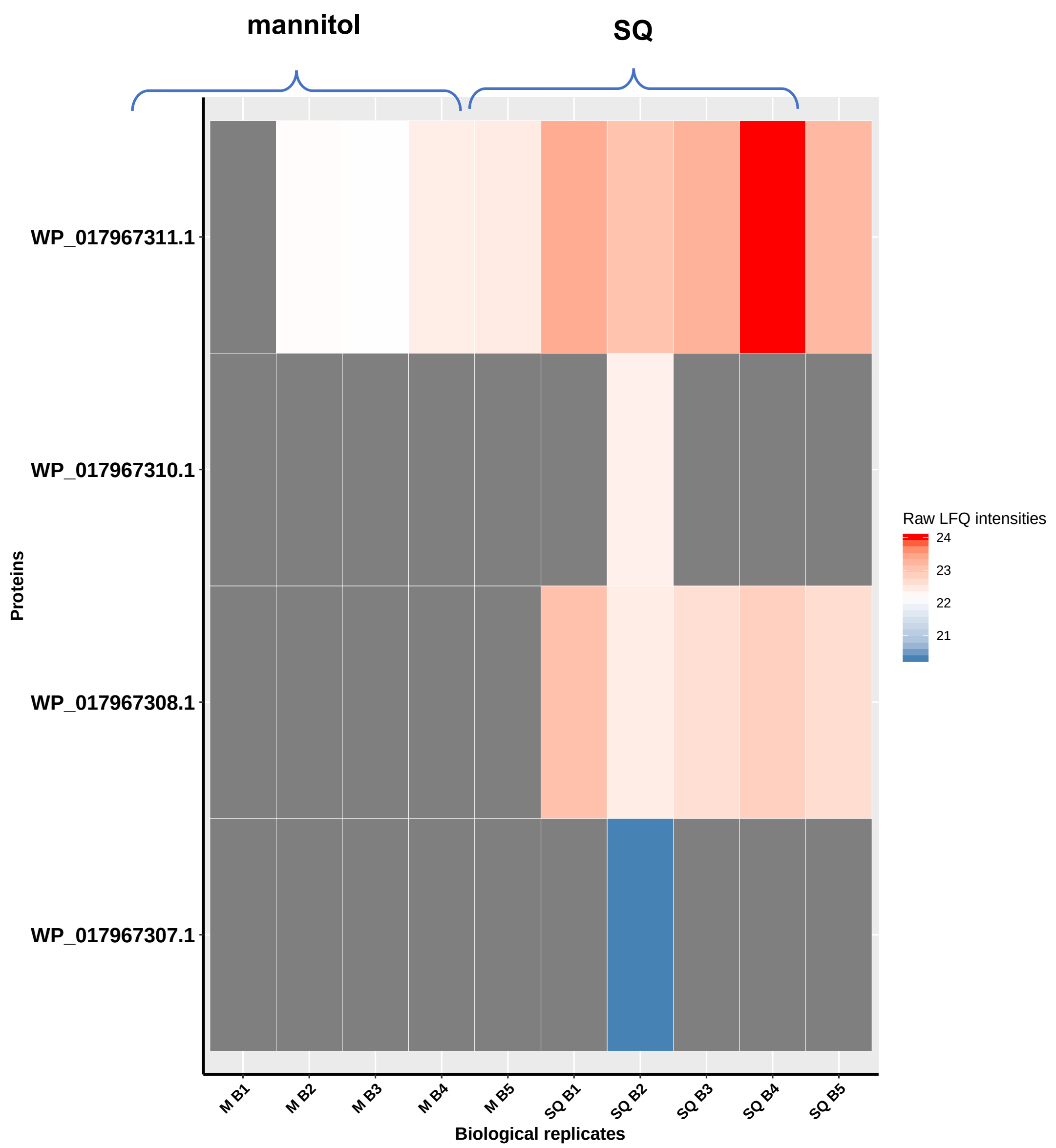


**Figure S7.** Heatmap of proteomics LFQ intensity data showing putative sulfo-ED operon protein observed across individual growth experiments of *Rl*-SRDI565.

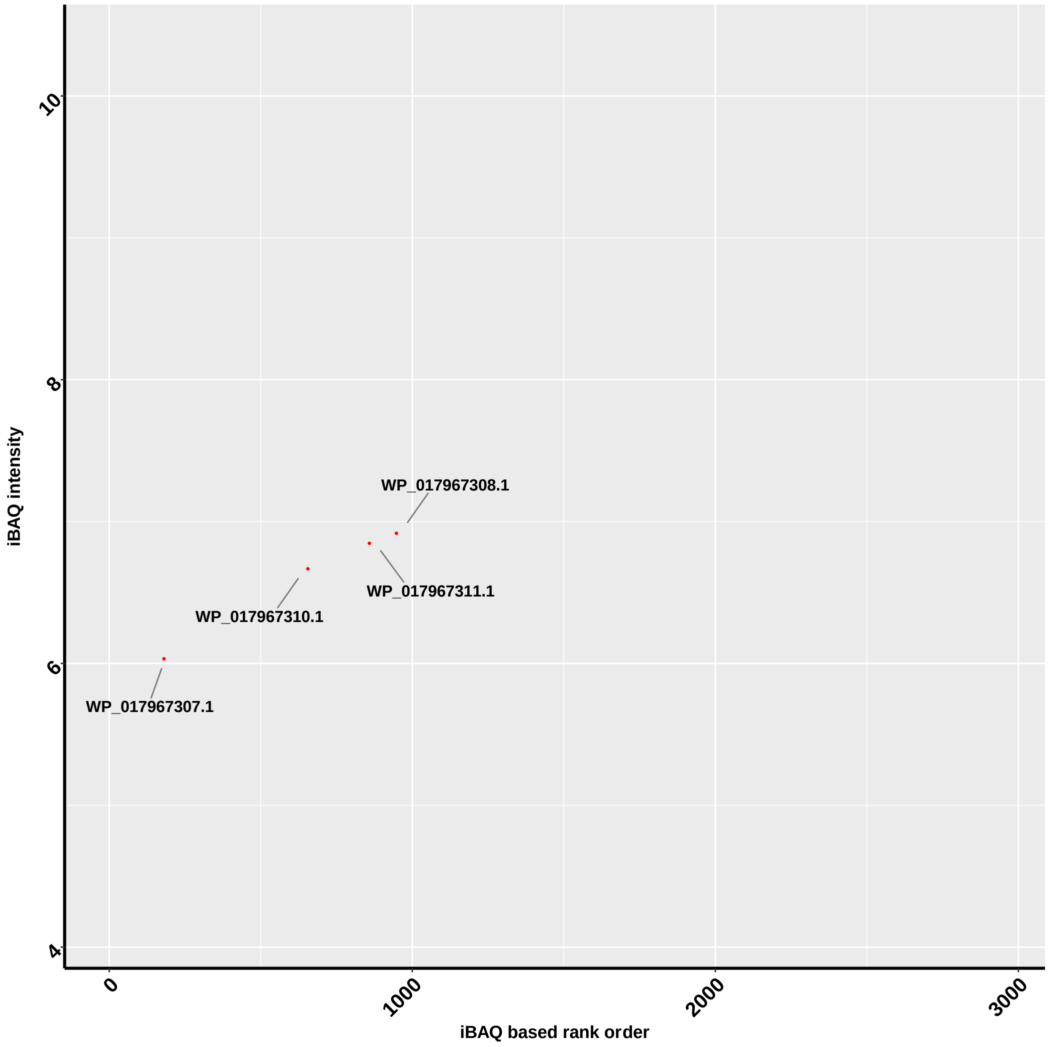


**Figure S8.** Rank plot of proteins detected from Rl-SRDI565 from SQ media determined by LC-MS/MS using intensity-based absolute quantification (iBAQ). The four proteins indicated with a red circle are proteins from the putative sulfo-ED pathway. iBAQ based analysis supports these proteins are found at low abundance during growth of SQ.

| Proposed Mutations: | |
|---|---|
| **Cluster #1:**<br>**SERp Score: 4.45** | Residues 375 - 376: KK |
| · K 375 => A<br>· K 376 => A | |
| **Cluster #2:**<br>**SERp Score: 4.4** | Residues 330 - 333: AEEA |
| · E 331 => A<br>· E 332 => A | |
| **Cluster #3:**<br>**SERp Score: 3.8** | Residues 35 - 36: EE |
| · E 35 => A<br>· E 36 => A | |
| **Cluster #4:**<br>**SERp Score: 3.47** | Residues 445 - 446: QK |
| · Q 445 => A<br>· K 446 => A | |

**Amino acid Sequence: SERp mutant K375/376A**

MKLEQTETGFILTFRGRTILEHSAAAPAIFAGHGEERMDMYRGNFEIEDYVIERRAFAHTEIEHNRVFLSDGPGRPAR
LVLNIDGGTIGFEALDTSLNRLWMRVPADHDEHVWGGGEQMSYLDMRGRRFPLWTSEPGVGRDRTSEITFKSNV
KDRAGGDYFNTNYPQPTYVSSARYALHVETTAYSVFDFRRKTFHEIEVWAIPSRIELFAAPTFVELVEVLSDRFGRQ
PPLPDWVYNGAIIGLKDGENSFSRLQLMRDAGVEVSALWCEDWVGLRQTSFGARLFWDWQANEDRYPGLRQRIT
ELNGDGIRFLGYVNPYLCVDGPLFEIAEAEGYFATDESGKTALVDFGEFDCGVVDFTNPAAAEWFAEDIIGAAMLDY
GLSGWMADFGEYLPIDVHLANGIDAKLMHNQWPTLWAEVNAKAVASRGQTGEALFFMRAGFTGVQKHCPLLWGG
DQSVDFSRHDGLVTVICAALSSGLLGNAYHHSDIGGYTSLFGNVRTPELLMRWAEMAAFTPVMRSHEGNRPRENV
QIDQDAQVLSHFARMTRIYRHLAPYLKSLSDEAVARGLPVQRPLFLHHQDDRETYAIQDSYLYGADMLVAPVWEAA
QVDRVVYLPQGERWVHAWTGETFAGGEKVRIAAPLGQPPVFYRAGCADEALFAGLRGL*

**Figure S9.** (Top) Surface entropy reduction analysis of Rl-SRDI565 SQase. (Bottom) Amino acid sequence for the mutant K375A/376A (changes underlined), termed *Rl*SQase*.

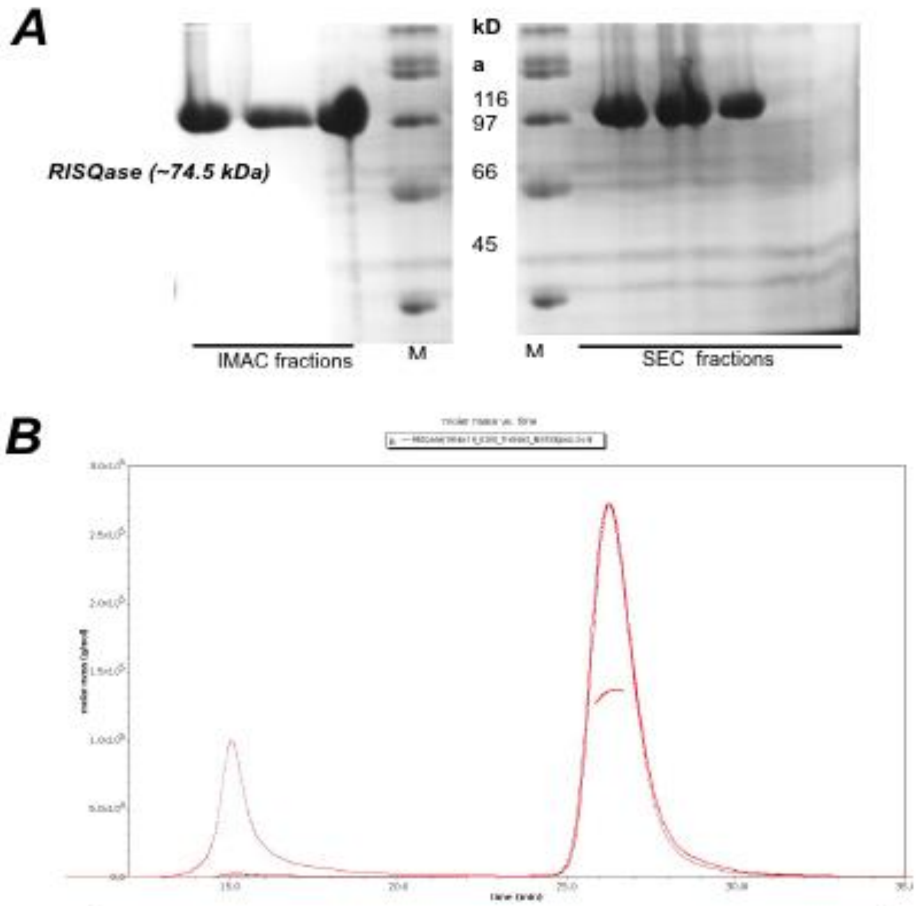


| Peak | from (min) | to (min) | Amount (µg) | MW (kDa) |
|---|---|---|---|---|
| Major Peak (99%) | 24.00 | 32.00 | 353 | 134 |
| Minor peak | 21.60 | 24.20 | 3.5 | 3800 |
| Total | | | 356.5 | |

**Figure S10.** Purification and characterization of *Rl*SQase*. A) SDS-PAGE analysis of purified *Rl*SQase* after immobilized metal-ion affinity chromatography (left) and size exclusion chromatography (right) (expected MW: 74,500 Da). B) SEC-MALS plot reveals the oligomeric state of *Rl*SQase*in solution. Molar mass plots show the LS trace as a solid red line, the RI trace (concentration measurement) as a dashed red line, and the UV trace as a dotted red line. These are normalised to the largest peak giving an estimated mass of 134 kDa, which corresponds to a dimer..

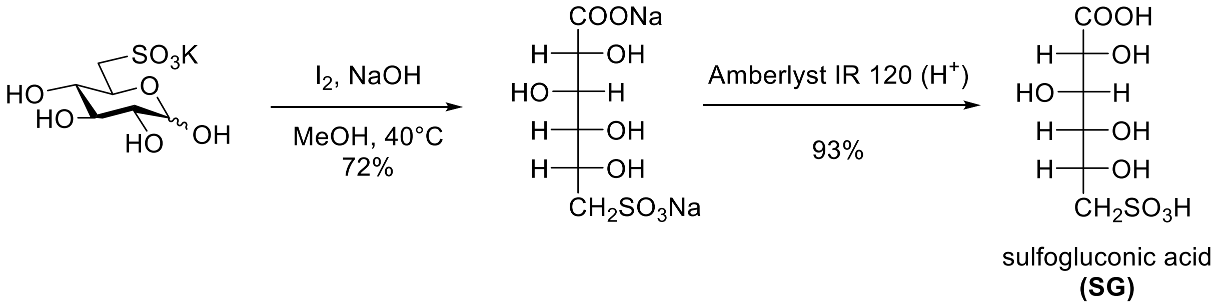


**Figure S11.** Chemical synthesis of sulfogluconic acid (SG).

$^{1}H$ NMR

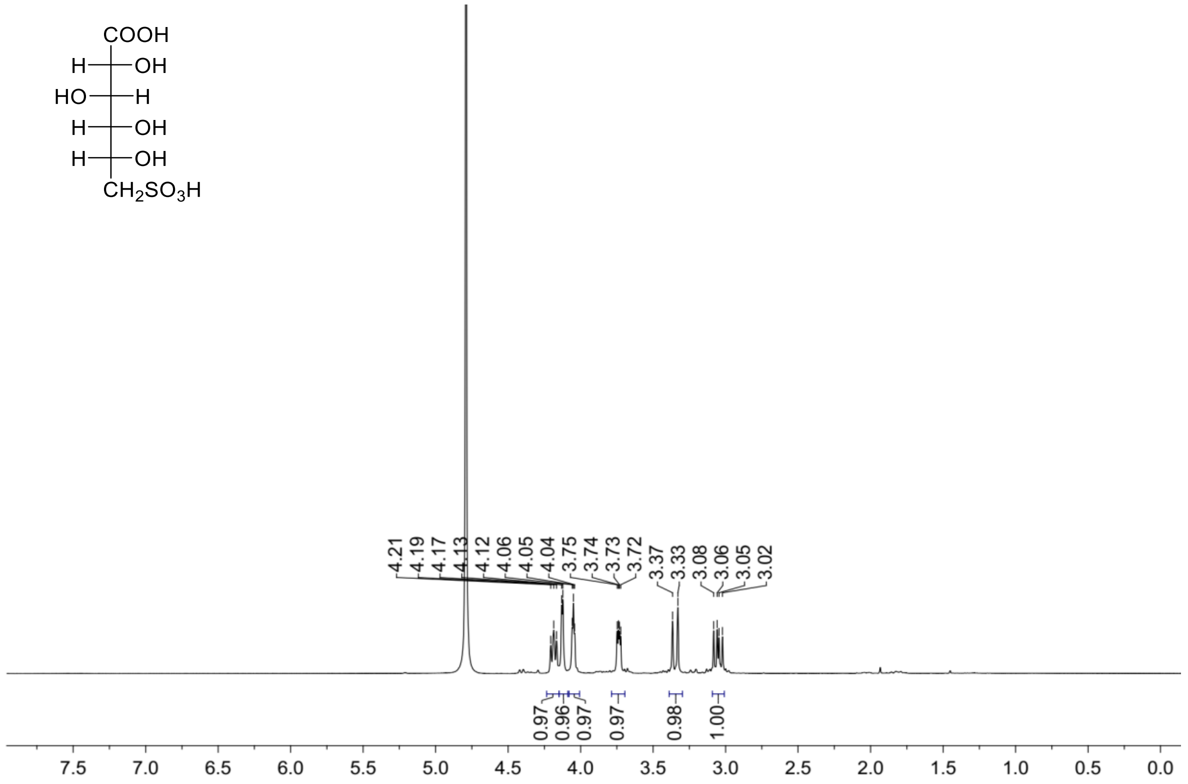


$^{13}C\{^{1}H\}$ NMR

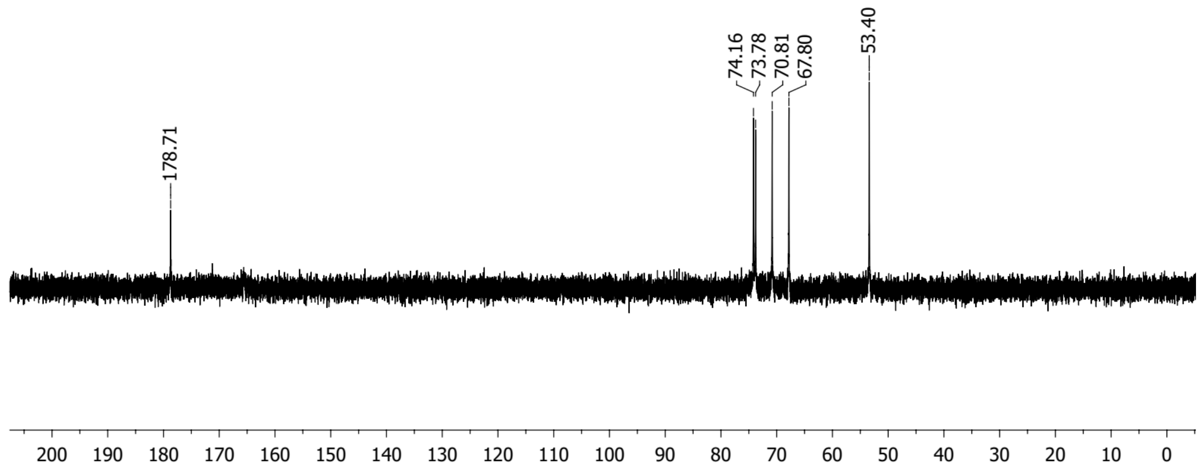


**Figure S12.** $^{1}H$ and $^{13}C\{^{1}H\}$ NMR spectra for 6-deoxy-6-sulfogluconic acid (SG).

**a)**

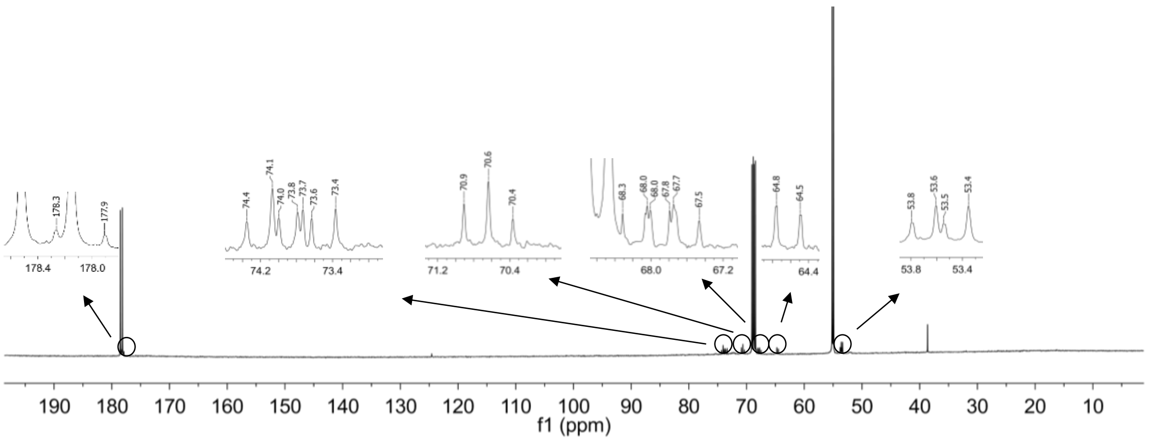


### b) Data for $^{13}C_6$-SG

| $^{13}$C chemical shift (δ ppm) | $^1J(^{13}C\text{-}^{13}C)$ coupling (Hz) | assignment |
|---|---|---|
| 178.1 | $J_{1,2} = 53.2$ | C1, d |
| 70.6 | $J_{2,3} \approx J_{3,4} \approx 40.9$ | C3, t |
| 74.1 | $J_{3,4} \approx J_{4,5} \approx 42.5$ | C4, t |
| 73.7 | $J_{1,2} = 53.2$ | C2, dd |
| | $J_{2,3} = 40.9$ | |
| 67.8 | $J_{4,5} = 42.5$ | C5, dd |
| | $J_{5,6} = 38.3$ | |
| 53.5 | $J_{5,6} = 38.3$ | C6, d |

### c) Data for $^{13}C_3$-DHPS

| $^{13}$C chemical shift (δ ppm) | $^1J(^{13}C\text{-}^{13}C)$ coupling (Hz) | assignment |
|---|---|---|
| 53.7 | $J_{1,2} = 37.5$ | C3, d |
| 68.1 | $J_{2,3} = 41.1$ | C2, dd |
| | $J_{1,2} = 37.5$ | |
| 64.6 | $J_{2,3} = 41.1$ | C1, d |

**Figure S13. $^{13}$C NMR (151 MHz) analysis of culture media from growth of *Rl*-SRDI565 on $^{13}C_6$-SQ using a cryoprobe for enhanced sensitivity.** a) $^{13}C\{^1H\}$ NMR spectrum with expansions of minor signals. b) Tabulated $^{13}$C NMR data assigned to $^{13}C_6$-sulfogluconate. C) Tabulated $^{13}$C NMR data assigned to $^{13}C_3$-dihydroxypropanesulfonate. All samples contain 10% $D_2O$, added to allow frequency lock.

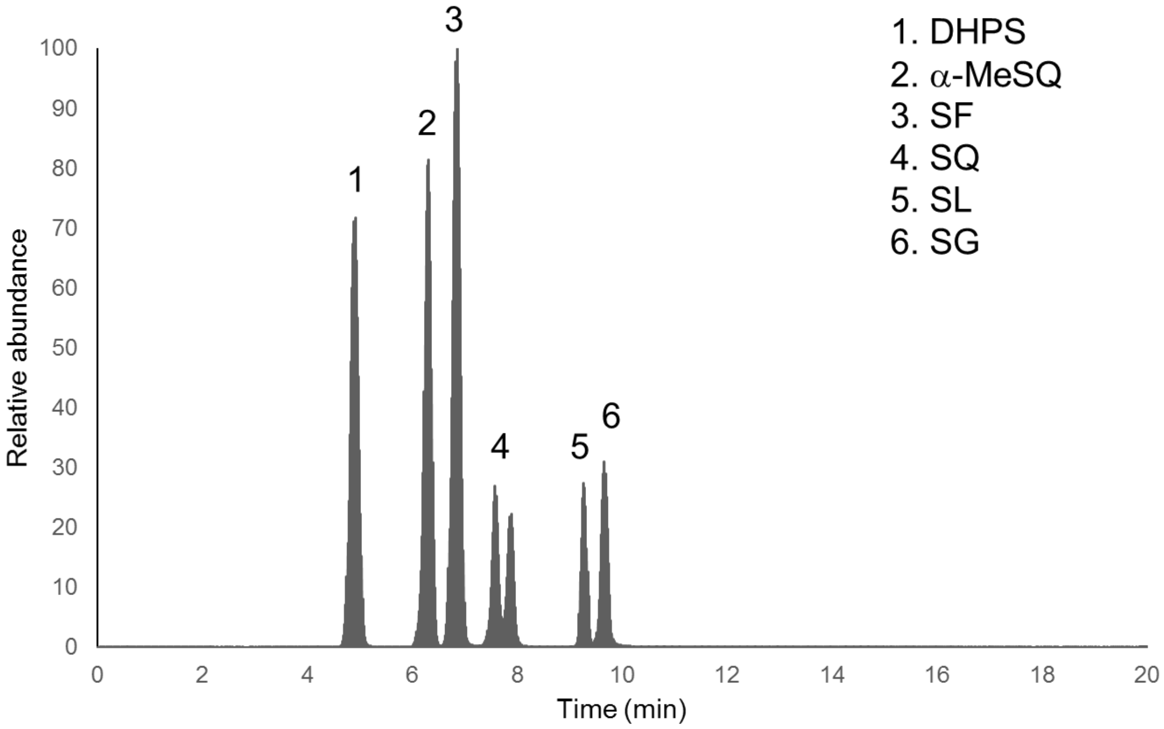


**Figure S14.** Total ion chromatogram of mixture of authentic sulfonate metabolites and internal standard (α-MeSQ) at 0.2 mM.

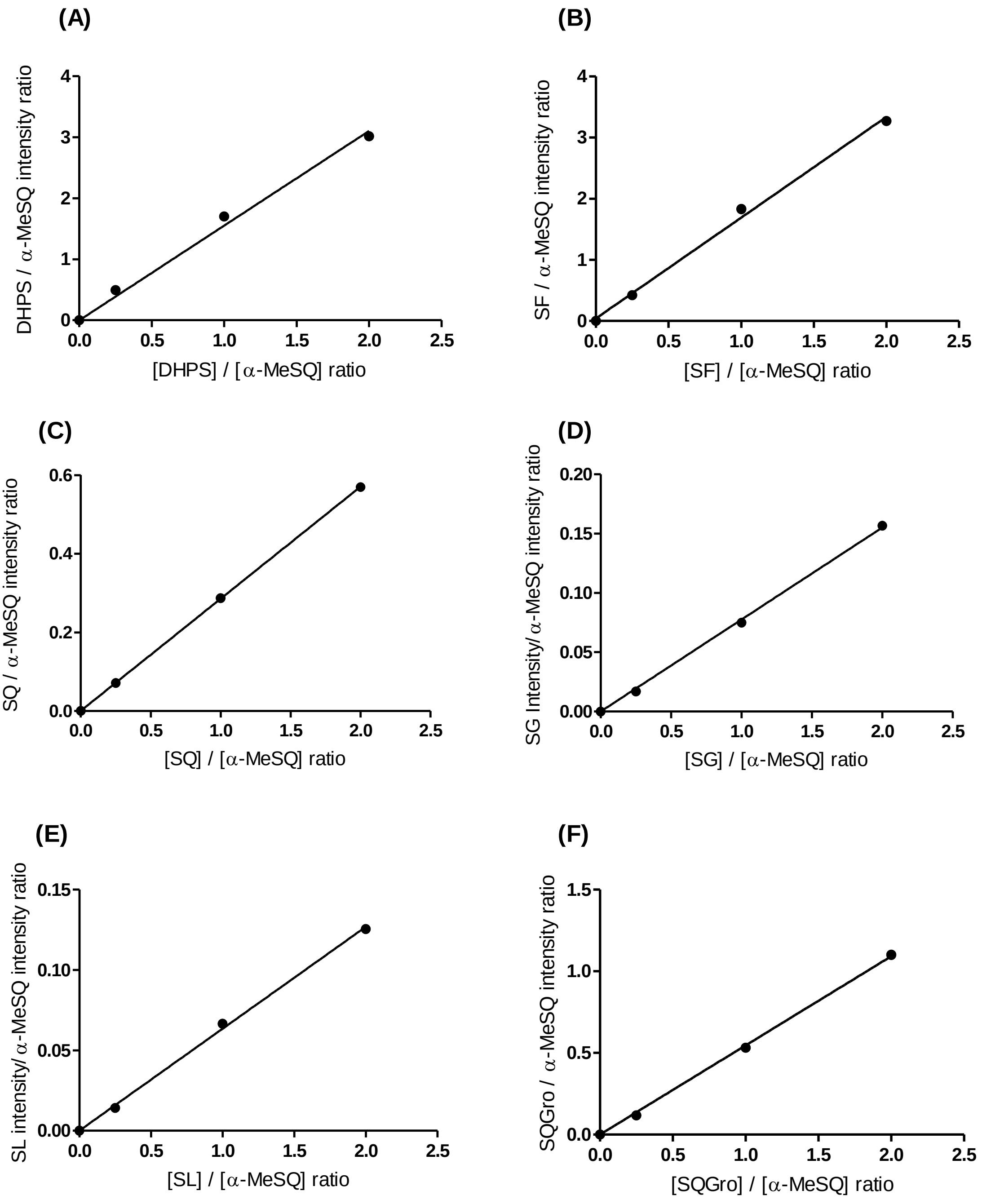


**Figure S15.** LC/MS-MS calibration curves for authentic, chemically synthesized DHPS, SF, SQ, SG, SL, and SQGro acquired using selected reaction monitoring (SRM) mode and normalized with respect to the internal standard, α-MeSQ.
